# Salt-dependent non-catalytic allostery of human USP14-regulated 26S proteasome

**DOI:** 10.1101/2024.11.07.622408

**Authors:** Shitao Zou, Shuwen Zhang, Lihong Zhao, Youdong Mao

## Abstract

USP14, a deubiquitylating enzyme, regulates 26S proteasome function both catalytically and non-catalytically through multiple checkpoints. How USP14 non-catalytically regulates the proteasome activity remains elusive. Here, we combined genetic engineering and cryo-EM to disentangle how USP14 non-catalytically regulates proteasome activity in a salt-dependent manner. By solving 16 high-resolution cryo-EM structures of substrate-engaged human 26S proteasome complexed with a catalytically deficient mutant USP14, we demonstrate that USP14’s non-catalytic activity alone can induce parallel pathways of proteasome state transitions, leading to opposing substrate fates. The USP14 mutant allosterically reprograms the AAA-ATPase motor, inducing novel substrate-engaged conformations and filling major gaps in understanding asymmetric ATP-hydrolysis cycling around the ATPase ring. Time-resolved cryo-EM and functional analysis reveal that sodium or potassium promotes substrate-engaged pathways and suppresses USP14 activity for enhanced degradation, uncovering another layer of complexity in proteasome regulation.

---

Ubiquitin-mediated proteolysis plays a critical role in various cellular processes across eukaryotic organisms^1-3^. Proteins tagged with ubiquitin are degraded by the 26S proteasome^4-7^, a complex machine comprising a 700-kDa barrel-shaped 20S core particle (CP) and one or two 900-kDa 19S regulatory particles (RPs). At the heart of the RP is a ring-shaped motor module formed by six AAA (ATPase associated with diverse cellular activities)-ATPase subunits (RPT1-RPT6), which utilize the chemical energy of ATP hydrolysis to unfold substrates and thread them into the proteolytic chamber of CP^8-11^. The RP process substrates through several sequential intermediate steps, including ubiquitin recognition by the ubiquitin receptor subunits RPN1, RPN10 or RPN13^12^, substrate engagement with the central channel of AAA-ATPase motor via a flexible initiation region^13^, ATP-dependent deubiquitylation^14-16^, and substrate unfolding and translocation into the CP by the AAA-ATPase motor^17,18^.

The stoichiometric deubiquitylating enzyme (DUB) subunit RPN11 is essential for the proteasome function and catalyzes removal of ubiquitin in an ATP-dependent manner to facilitate substrate degradation^14-16^. However, ubiquitylated substrates can be rescued from degradation through early removal of conjugated ubiquitin by other DUBs. Two such DUBs, USP14 (ubiquitin-specific protease 14, also known as Ubp6 in yeast) and UCH37 (also known as UCHL5), reversibly associate with the proteasome to fine-tune its activity^19,20^. USP14 is pivotal in numerous cellular processes and is a promising therapeutic target for diseases like cancer and neurodegeneration^21-23^. Its N-terminal ubiquitin-like (UBL) domain binds the toroidal domain of proteasome subunit RPN1^24^. Its C-terminal palm-like USP domain harbors a catalytic cysteine in the binding pocket for the ubiquitin’s C-terminus. The blocking loop 1 (BL1) of USP14 binds the oligonucleotide- or oligosaccharide-binding (OB) domain of the proteasomal ATPase subunit RPT1, opening up its catalytic pocket for activation^25-29^ and promoting the *en bloc* removal of ubiquitin chains from substrates^30-32^.

USP14 suppresses proteasomal degradation and inhibits the DUB activity of RPN11 both catalytically and non-catalytically^20,30,31,33,34^. The non-catalytic effect is inherently coupled with USP14 activation on the proteasome^28^. In certain in vivo contexts, the non-catalytic effect can be more influential, affecting cellular protein turnover independently of deubiquitylation^30,34,35^. In the absence of USP14, cryo-EM reconstructions have captured key proteasome conformations during substrate processing, including states E_A_ (ubiquitin recognition), E_B_ (RPN11-catalysed deubiquitylation), E_C_ (initiation of substrate translocation) and E_D_ (processive translocation)^9,36^. Recent time-resolved cryo-EM studies revealed that USP14 introduces multiple regulatory checkpoints on the proteasome by inducing both substrate-engaged and substrate-inhibited pathways of proteasome state transition, bypassing the intermediate conformational states E_B_ and E_C_ ^29^. On the substrate-inhibited pathway, substrate entry into the central channel of the AAA-ATPases is sterically occluded by RPN11, allowing USP14 to rescue substrates. This dual-pathway model provides a framework for understanding how USP14 allosterically regulates the proteasome during substrate degradation.

However, several outstanding problems persist. First, all previously identified conformations of the USP14-proteasome complex correspond to post-deubiquitylation states, where the USP14-bound ubiquitin has been separated from substrate. The pre-deubiquitylation states remain unidentified. Second, the catalytic and non-catalytic effects of USP14 have been entangled and observed in a holistic fashion in the earlier structural studies^29^, making it difficult to precisely determine the mechanism of the non-catalytic effect. Third, proteasome substrates display a wide range of sensitivities to non-catalytic regulation or inhibition by USP14^30^, implicating a more complex regulatory role in USP14 activity that is not yet well understood. How the physiological electrolytes such as sodium and potassium influence USP14-mediated proteasome regulation remains unknown.

In this study, we aimed to address these problems by mutating the catalytic cysteine at residue 114 of USP14 to alanine (referred to as USP14^C114A^) and determined 16 cryo-EM structures of the USP14^C114A^-proteasome complex in the presence of polyubiquitylated substrates with RPN11 inhibited. These structural snapshots reveal previously unidentified pre-deubiquitylation states of the substrate-engaged USP14-proteasome, including key intermediate conformations that occur prior to the USP14^C114A^-induced bifurcation of state-transition pathways. Importantly, we captured a series of previously missing conformations of substrate-engaged proteasome and salt-modulated variation of state-transition pathways, offering novel insights into how USP14 non-catalytically regulates the proteasome in a salt-dependent manner.

## Results

### Pre-deubiquitylation conformers of USP14^C114A^-proteasome

To prepare the human proteasome in complex with the catalytically deficient USP14^C114A^ mutant, we incubated the USP14-free proteasome with purified USP14^C114A^ in a buffer containing the RPN11 inhibitor ortho-phenanthroline (*o*PA)^15^. To better correlate structural analysis with functional assays, we genetically engineered a new model substrate, designated NtSic1^PY^-cp8sGFP, by fusing the N-terminal region of Sic1^PY^ with a superfolder GFP mutant (see Methods). The substrate was polyubiquitylated *in vitro* (Extended Data Fig. 1a-c) and then mixed with the reconstituted USP14^C114A^-proteasome complex for single-particle cryo-EM analysis. By using a hierarchical 3D classification approach (Extended Data Fig. 2), we determined 16 distinct conformational states of the USP14^C114A^-proteasome complex in the act of substrate processing. The nominal resolutions for the USP14^C114A^-RP subcomplex ranged from 3.6 to 4.5 Å, including four E_A_-like, two S_D_-like (substrate-inhibited) and ten E_D_-like (substrate-engaged) states (Fig. 1a, Extended Data Figs. 3-7, and Tables 1 and 2). In the four E_A_-like states, the catalytic zinc ion at the active site of RPN11 is visible, but it is absent in E_D_-like and S_D_-like states due to *o*PA treatment, which chelates and sequesters the zinc ion (Extended Data Fig. 5q). Notably, despite exhaustive 3D classification, we did not observe states E_B_ and E_C_ that were originally identified in the USP14-free proteasome, consistent with previous studies on the 26S proteasome complexed with wild-type USP14 (USP14^WT^)^29^.

**Table 1:** Cryo-EM data collection, refinement and validation statistics.

|  | E<sub>A1</sub> | E<sub>A1</sub><sup>USP14ca</sup> | E<sub>A2.1</sub> | E<sub>A2.1</sub><sup>USP14ca</sup> | S<sub>B</sub><sup>USP14ca</sup> | S<sub>D4.0</sub><sup>USP14ca</sup> | E<sub>D0.0</sub><sup>USP14ca</sup> | E<sub>D0.1</sub><sup>USP14ca</sup> |
| --- | --- | --- | --- | --- | --- | --- | --- | --- |
| EMDB ID | 61888 | 61889 | 61890 | 61891 | 61892 | 61893 | 61894 | 61895 |
| PDB ID | 9JY5 | 9JY6 | 9JY7 | 9JY8 | 9JY9 | 9JYA | 9JYB | 9JYC |
| **Data collection and processing** |  |  |  |  |  |  |  |  |
| Magnification | 105,000 | 105,000 | 105,000 | 105,000 | 105,000 | 105,000 | 105,000 | 105,000 |
| Voltage (kV) | 300 | 300 | 300 | 300 | 300 | 300 | 300 | 300 |
| Electron exposure (e<sup>-</sup>/Å<sup>2</sup>) | 50 | 50 | 50 | 50 | 50 | 50 | 50 | 50 |
| Defocus range (μm) | -0.4 to -4.0 | -0.4 to -4.0 | -0.4 to -4.0 | -0.4 to -4.0 | -0.4 to -4.0 | -0.4 to -4.0 | -0.4 to -4.0 | -0.4 to -4.0 |
| Pixel size (Å) | 0.668 | 0.668 | 0.668 | 0.668 | 0.668 | 0.668 | 0.668 | 0.668 |
| Symmetry imposed | C1 | C1 | C1 | C1 | C1 | C1 | C1 | C1 |
| Initial particle images (no.) | 1,856,819 | 1,856,819 | 1,856,819 | 1,856,819 | 1,856,819 | 1,856,819 | 1,856,819 | 1,856,819 |
| Final particle images (no.) | 54,701 | 15,531 | 14,827 | 9,208 | 20,096 | 12,157 | 49,551 | 20,545 |
| Map resolution (Å) | 3.6 (RP) | 4.1 (RP) | 4.2 (RP) | 4.4 (RP) | 4.5 (RP) | 4.3 (RP) | 3.6 (RP) | 4.0 (RP) |
|  | 3.0 (CP) | 3.4 (CP) | 3.4 (CP) | 3.5 (CP) | 3.3 (CP) | 3.4 (CP) | 2.9 (CP) | 3.1 (CP) |
| FSC threshold | 0.143 | 0.143 | 0.143 | 0.143 | 0.143 | 0.143 | 0.143 | 0.143 |
| Map resolution range (Å) | 2.7-5.0 | 2.7-7.0 | 2.7-6.0 | 2.7-8.0 | 2.7-8.0 | 2.7-7.0 | 2.7-5.0 | 2.7-6.0 |
| **Refinement** |  |  |  |  |  |  |  |  |
| Initial model used (PDB code) | 7W37 | 7W37 | 7W39 | 7W39 | 7W3I | 7W3J | 7W3C | 7W3C |
| Model resolution (Å) | 3.3 | 3.8 | 3.6 | 3.9 | 3.4 | 3.7 | 3.2 | 3.4 |
| FSC threshold | 0.5 | 0.5 | 0.5 | 0.5 | 0.5 | 0.5 | 0.5 | 0.5 |
| Map sharpening *B* factor (Å<sup>2</sup>) | -50 | -20 | -20 | -20 | -20 | -20 | -50 | -30 |
| **Model composition** |  |  |  |  |  |  |  |  |
| Non-hydrogen atoms | 105,015 | 109,256 | 105,117 | 108,488 | 110,329 | 110,369 | 111,191 | 111,089 |
| Protein residues | 13,388 | 13,918 | 13,375 | 13,794 | 14,028 | 14,039 | 14,130 | 14,119 |
| Ligands | 12 | 7 | 7 | 7 | 6 | 6 | 9 | 6 |
| ***B* factors (Å<sup>2</sup>)** |  |  |  |  |  |  |  |  |
| Protein | 84.12 | 105.69 | 135.96 | 114.47 | 124.83 | 104.63 | 87.27 | 104.16 |
| Ligand | 70.48 | 84.41 | 151.18 | 122.86 | 110.87 | 114.19 | 83.34 | 117.71 |
| **R.m.s. deviations** |  |  |  |  |  |  |  |  |
| Bond lengths (Å) | 0.003 | 0.003 | 0.005 | 0.003 | 0.005 | 0.004 | 0.003 | 0.005 |
| Bond angles (°) | 0.666 | 0.737 | 0.784 | 0.786 | 0.789 | 0.788 | 0.693 | 0.777 |
| **Validation** |  |  |  |  |  |  |  |  |
| MolProbity score | 1.64 | 1.68 | 1.72 | 1.70 | 1.80 | 1.78 | 1.69 | 1.78 |
| Clashscore | 6.11 | 6.31 | 6.94 | 7.14 | 7.63 | 7.33 | 6.47 | 7.78 |
| Poor rotamers (%) | 0.42 | 0.15 | 0.19 | 0.16 | 0.36 | 0.26 | 0.64 | 0.64 |
| **Ramachandran plot** |  |  |  |  |  |  |  |  |
| Favored (%) | 95.63 | 95.22 | 95.21 | 95.52 | 94.46 | 94.44 | 95.22 | 94.80 |
| Allowed (%) | 4.29 | 4.73 | 4.71 | 4.38 | 5.40 | 5.41 | 4.62 | 5.06 |
| Disallowed (%) | 0.08 | 0.05 | 0.08 | 0.10 | 0.14 | 0.14 | 0.16 | 0.14 |

|  | E<sub>D1.0</sub><sup>USP14ca</sup> | E<sub>D1.1</sub><sup>USP14ca</sup> | E<sub>D2.0</sub><sup>USP14ca</sup> | E<sub>D2.1</sub><sup>USP14ca</sup> | E<sub>D2.2</sub><sup>USP14ca</sup> | E<sub>D3</sub><sup>USP14ca</sup> | E<sub>D4.0</sub><sup>USP14ca</sup> | E<sub>D4.1</sub><sup>USP14ca</sup> |
| --- | --- | --- | --- | --- | --- | --- | --- | --- |
| EMDB ID | 61896 | 61897 | 61898 | 61899 | 61900 | 61901 | 61902 | 61903 |
| PDB ID | 9JYD | 9JYE | 9JYF | 9JYG | 9JYH | 9JYI | 9JYJ | 9JYK |
| **Data collection and processing** |  |  |  |  |  |  |  |  |
| Magnification | 105,000 | 105,000 | 105,000 | 105,000 | 105,000 | 105,000 | 105,000 | 105,000 |
| Voltage (kV) | 300 | 300 | 300 | 300 | 300 | 300 | 300 | 300 |
| Electron exposure (e<sup>-</sup>/Å<sup>2</sup>) | 50 | 50 | 50 | 50 | 50 | 50 | 50 | 50 |
| Defocus range (μm) | -0.4 to -4.0 | -0.4 to -4.0 | -0.4 to -4.0 | -0.4 to -4.0 | -0.4 to -4.0 | -0.4 to -4.0 | -0.4 to -4.0 | -0.4 to -4.0 |
| Pixel size (Å) | 0.668 | 0.668 | 0.668 | 0.668 | 0.668 | 0.668 | 0.668 | 0.668 |
| Symmetry imposed | C1 | C1 | C1 | C1 | C1 | C1 | C1 | C1 |
| Initial particle images (no.) | 1,856,819 | 1,856,819 | 1,856,819 | 1,856,819 | 1,856,819 | 1,856,819 | 1,856,819 | 1,856,819 |
| Final particle images (no.) | 14,661 | 24,812 | 50,917 | 92,349 | 36,029 | 15,976 | 47,644 | 35,525 |
| Map resolution (Å) | 4.2 (RP) | 4.0 (RP) | 3.8 (RP) | 3.6 (RP) | 3.8 (RP) | 4.4 (RP) | 3.7 (RP) | 3.9 (RP) |
|  | 3.4 (CP) | 3.2 (CP) | 3.0 (CP) | 2.8 (CP) | 3.0 (CP) | 3.2 (CP) | 3.0 (CP) | 3.0 (CP) |
| FSC threshold | 0.143 | 0.143 | 0.143 | 0.143 | 0.143 | 0.143 | 0.143 | 0.143 |
| Map resolution range (Å) | 2.7-8.0 | 2.7-6.0 | 2.7-5.0 | 2.7-5.0 | 2.7-5.0 | 2.7-6.0 | 2.7-5.0 | 2.7-5.0 |
| **Refinement** |  |  |  |  |  |  |  |  |
| Initial model used (PDB code) | 7W3F | 7W3F | 7W3G | 7W3H | 7W3H | 7W3A | 7W3A | 7W3A |
| Model resolution (Å) | 3.7 | 3.5 | 3.2 | 3.1 | 3.2 | 3.4 | 3.1 | 3.3 |
| FSC threshold | 0.5 | 0.5 | 0.5 | 0.5 | 0.5 | 0.5 | 0.5 | 0.5 |
| Map sharpening *B* factor (Å<sup>2</sup>) | -20 | -30 | -50 | -50 | -30 | -20 | -50 | -50 |
| Model composition |  |  |  |  |  |  |  |  |
| Non-hydrogen atoms | 111,008 | 110,989 | 111,041 | 111,168 | 110,989 | 111,145 | 111,139 | 110,214 |
| Protein residues | 14,109 | 14,108 | 14,111 | 14,117 | 14,105 | 14,123 | 14,119 | 14,030 |
| Ligands | 6 | 5 | 6 | 10 | 5 | 6 | 10 | 6 |
| *B* factors (Å<sup>2</sup>) |  |  |  |  |  |  |  |  |
| Protein | 116.90 | 110.20 | 93.85 | 89.13 | 91.85 | 114.11 | 84.22 | 100.57 |
| Ligand | 132.74 | 107.41 | 103.32 | 102.92 | 92.14 | 133.42 | 89.06 | 106.10 |
| R.m.s. deviations |  |  |  |  |  |  |  |  |
| Bond lengths (Å) | 0.004 | 0.004 | 0.003 | 0.004 | 0.003 | 0.005 | 0.003 | 0.003 |
| Bond angles (°) | 0.772 | 0.754 | 0.712 | 0.703 | 0.702 | 0.807 | 0.706 | 0.723 |
| Validation |  |  |  |  |  |  |  |  |
| MolProbity score | 1.74 | 1.75 | 1.72 | 1.70 | 1.71 | 1.77 | 1.70 | 1.73 |
| Clashscore | 6.95 | 7.17 | 6.74 | 6.42 | 6.71 | 7.00 | 6.59 | 6.99 |
| Poor Rotamers (%) | 0.17 | 0.55 | 0.53 | 0.44 | 0.49 | 0.23 | 0.46 | 0.55 |
| Ramachandran plot |  |  |  |  |  |  |  |  |
| Favored (%) | 94.88 | 94.89 | 95.03 | 95.04 | 95.05 | 94.43 | 95.10 | 95.01 |
| Allowed (%) | 5.00 | 4.97 | 4.84 | 4.78 | 4.79 | 5.44 | 4.78 | 4.83 |
| Disallowed (%) | 0.12 | 0.14 | 0.13 | 0.18 | 0.16 | 0.12 | 0.12 | 0.17 |

**Table 2:**
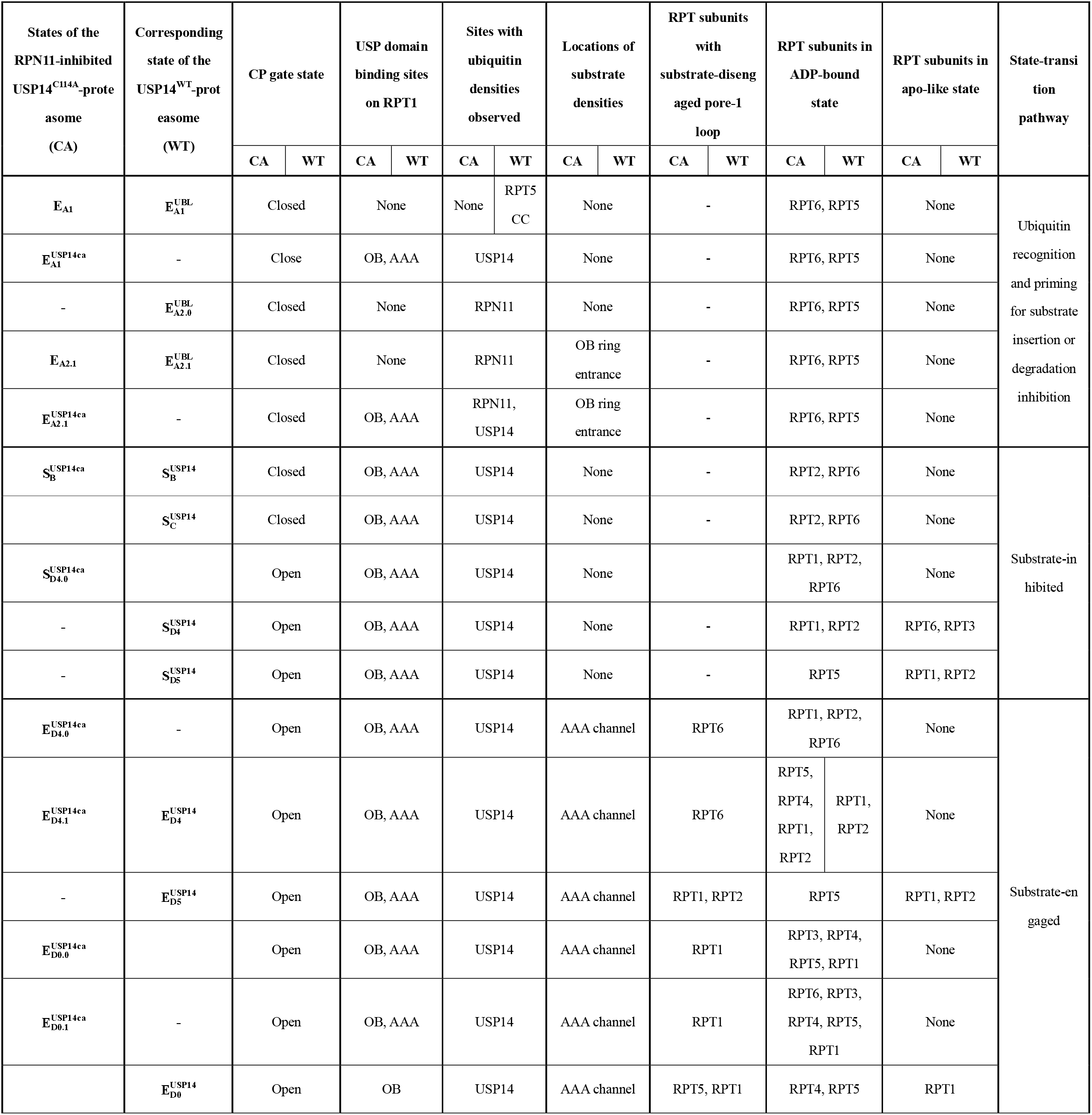
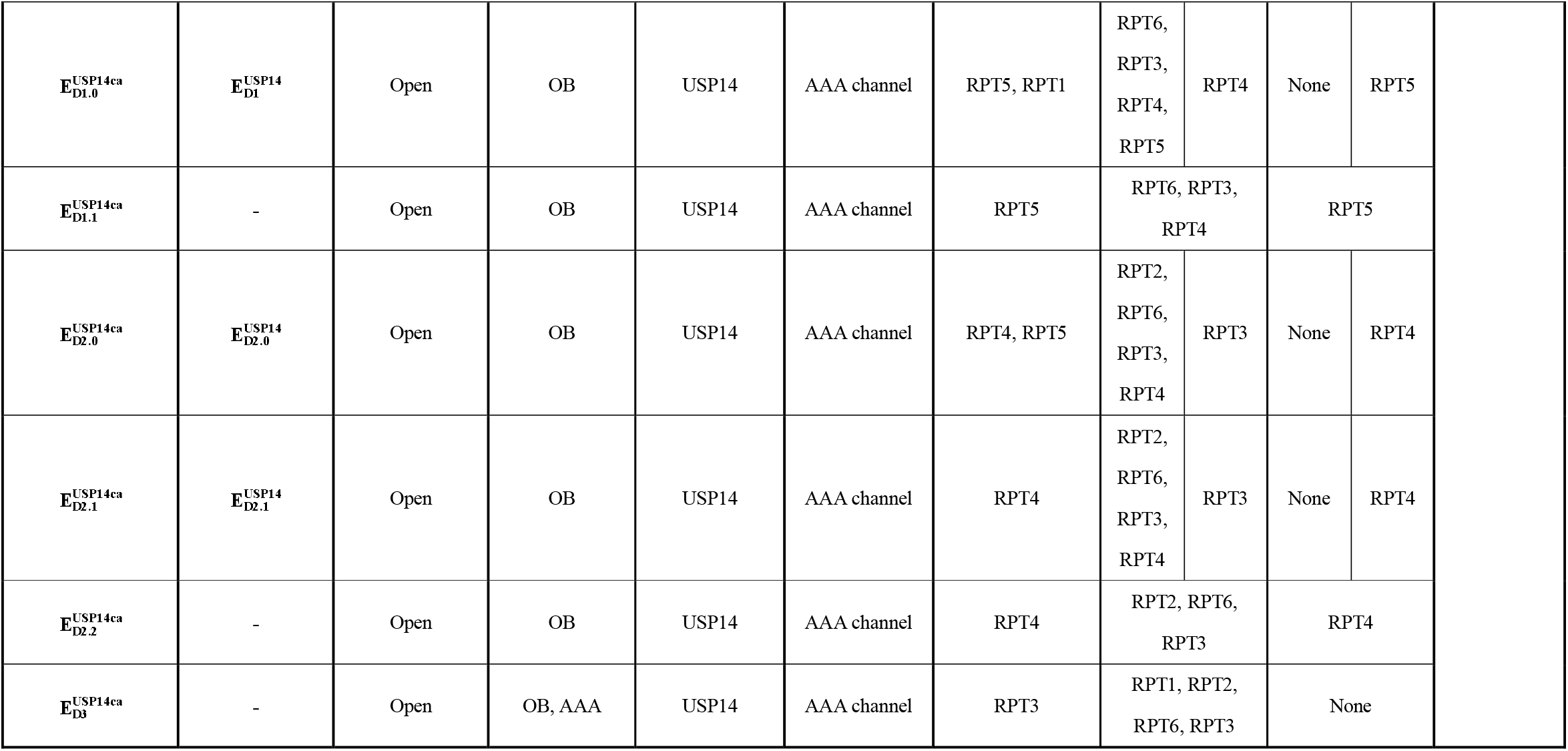
Summary of key structural features of the USP14-bound proteasome in different states. To provide a more comprehensive understanding of these structural features, the table includes previously reported conformational states of the USP14^WT^-proteasome^29^ (second column), regardless of whether they correspond to the states observed in this study (first column).

**Fig. 1:**
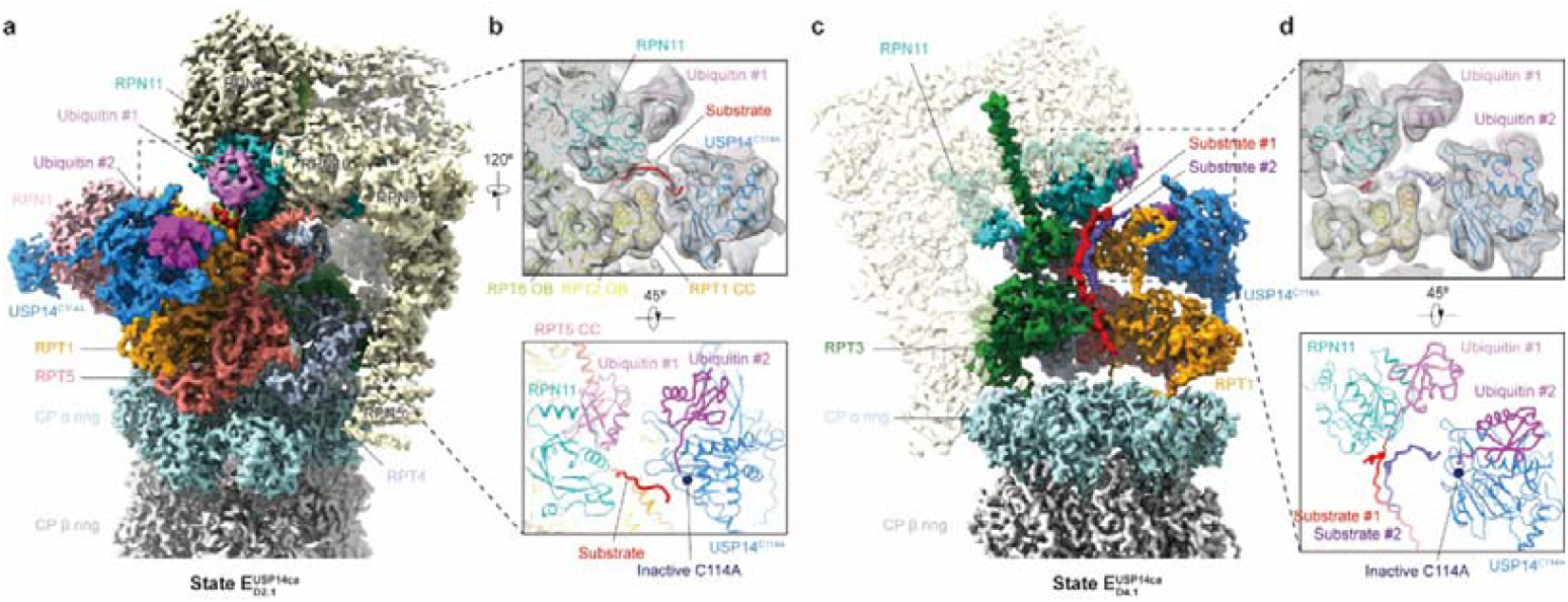
Cryo-EM structures of the substrate-engaged RPN11-inhibited USP14^C114A^-proteasome complex. **a**, Overview of the cryo-EM density map of the USP14^C114A^-proteasome in state 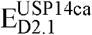. **b**, Close-up view of the substrate strand outside the OB ring in state 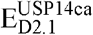. **c**, Cryo-EM density map of the substrate-engaged USP14^C114A^-proteasome in state 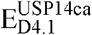. To better visualize the substrate density inside the central channel of the AAA-ATPase ring, the densities of RPN1, RPT2 and RPT6 are omitted, and all the RPN subunits, except for RPN11, are rendered transparent. **d**, Close-up view of the two substrate strands inside the OB ring in state 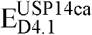. The upper images in panels **b** and **d** show the density map (transparent grey surface) low-pass filtered to 6 Å and superimposed with its atomic model, while the lower images display models of RPN11, USP14 and the ubiquitin-conjugated substrates. The isopeptide bond between the USP14-bound ubiquitin and the substrate is not built into the atomic model due to limited local resolution.

Two E_A_-like states, E_A1_ and E_A2.1_, lack visible densities for USP14, including the absence of UBL domain on RPN1, and exhibit RP conformations similar to those previously reported in states 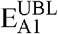and 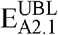 of the USP14^WT^-proteasome^29^, respectively (Extended Data Fig. 8k,m). In contrast to the absence of USP domain densities in any E_A_-like states of USP14^WT^-proteasome^29^, the other two E_A_-like states 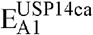 and 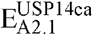 exhibit high-resolution densities of the ubiquitin-loaded USP^C114A^ domain attached onto the surface of the RPT1 OB domain, so do all E_D_-like and S_D_-like states (Fig. 1a and Extended Data Fig. 4a). In these two states, RPN1 is rotated aside about 60 Å around the AAA-ATPase ring compared to its position in states E_A1_ and E_A2.1_, which makes room for the USP^C114A^-RPT1 interaction (Fig. 2b). These conformations resemble the cryo-EM structure of ubiquitin aldehyde-bound USP14^WT^-proteasome in the absence of substrate^27^ (Extended Data Fig. 8b).

**Fig. 2:**
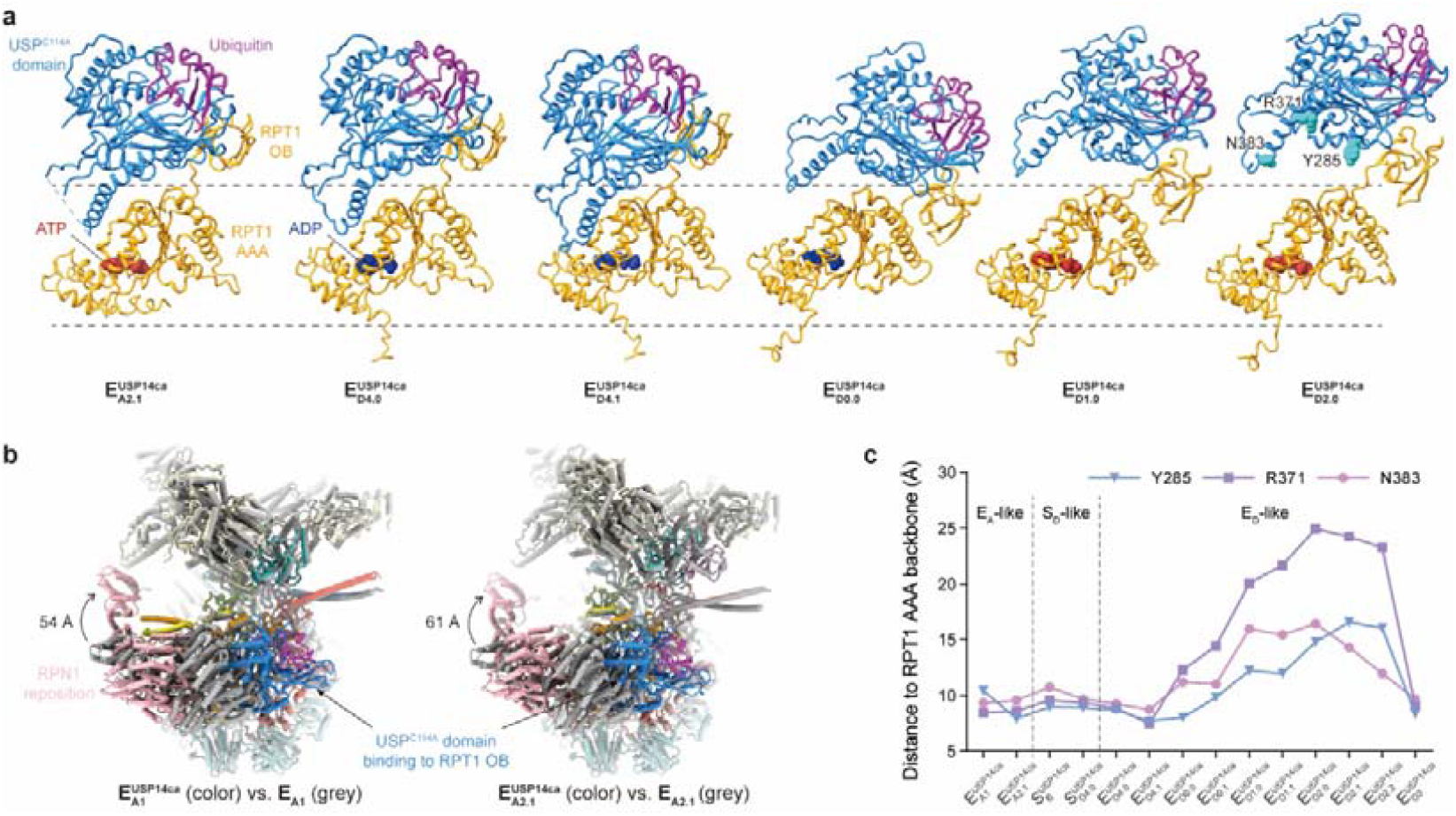
Dynamic USP^C114A^-RPT1 interactions. **a**, Side-by-side structural comparison of the USP^C114A^-ubiquitin-RPT1 subcomplex in six distinct states. The two dashed lines demarcate the RPT1 AAA domain used to align these models. ATP (red) and ADP (blue) in the nucleotide-binding pocket of RPT1 are highlighted in sphere representation. **b**, Structural comparison of the RP in the E_A_-like states before and after the binding of USP^C114A^ domain to RPT1. The models are superimposed after alignment of the CP. The models of parts of the CP and the USP14^C114A^ UBL domain are hidden for clarity. **c**, Changes in the USP^C114A^-AAA interface across different states, characterized by measuring the shortest distance between three USP14^C114A^ residues (Y285, R371 and N383) and the main chains of the RPT1 AAA domain.

The densities of substrate polypeptides are observed at the entrance of the OB ring underneath RPN11 in states E_A2.1_ and 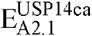, and inside the central channel of AAA-ATPase ring in all E_D_-like states (Extended Data Figs. 4a and 5a-p), but they are absent in the substrate-inhibited S_D_-like states (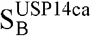 and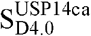). The CP gate is open in all E_D_-like states (Extended Data Fig. 5r), similar to previously reported substrate-translocating states^9,29,36^. Four E_D_-like states (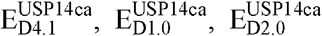 and 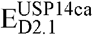) reproduce the previously reported E_D_-like states of USP14^WT^-proteasome in the overall RP conformation^29^ (Extended Data Figs. 8c and 9b,d,e, and Table 2). The other six E_D_-like states include a previously unidentified major conformation 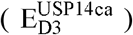 and several novel intermediate conformers characterizing a continuum of structural transitions in the AAA-ATPase motor (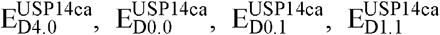 and 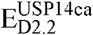). While state 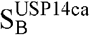 mostly reproduces state 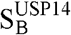 of the USP14^WT^-proteasome^29^ (Extended Data Fig. 9i), state 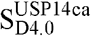 shows an open CP gate and RP conformation similar to 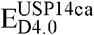 (Extended Data Figs. 5r and 7g).

### USP14 repositions substrate to override RPN11

To investigate how USP14^C114A^ regulates RPN11 activity, we performed focused 3D classification on a region encompassing both RPN11 and USP14^C114A^ using the cryo-EM particles of state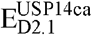, which resulted in an improved reconstruction of additional substrate densities in the RP (Fig. 1a,b). The substrate density extends from the proteasome central channel to the catalytic groove of RPN11, then turns towards the mutated C114A site of USP14, navigating through a narrow space between a helix of the RPT1 CC domain and the switching loop (SL) of USP14. USP14 must reorient the substrate in the vicinity, allowing the ubiquitin C-terminal segment to fully extend towards the catalytically active site of USP14 for swift cleavage. This spatial restriction of the substrate likely facilitates USP14 in capturing ubiquitin on the substrate compared to RPN11 as the substrate proceeds into the channel.

Unexpectedly, in state 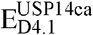, two twisted substrate strands are observed inside the OB ring with their associated ubiquitin captured by RPN11 and the USP^C114A^ domain (Fig. 1c,d and Extended Data Fig. 5p). While both substrate segments traverse the OB ring, only the one grasped by RPN11 is threaded into the central channel of the AAA-ATPase ring and reach the CP gate. These two substrate segments extend in divergent directions outside the OB ring (Fig. 1d), illustrating distinct paths for substrate segments processed by RPN11 and USP14, respectively. Although there are reported cases of the proteasome or other AAA proteins initiating substrate processing from internal loops^13,37,38^, the observed substrate polypeptide segments in the cryo-EM density map do not appear to be connected end-to-end. It is possible that these segments belong to entirely different substrates or distant regions of the same substrate, with the linking loop disordered and averaged out. Thus, USP14 may function in parallel with the proteasome, regardless of whether the proteasome is actively degrading a substrate. We hypothesize that the presence of two substrate segments in the OB ring might result from the optimal USP^C114A^-AAA interaction in state 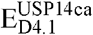 (Fig. 2c), which would stabilize and support the recognition of additional substrate segments.

As typically observed in states 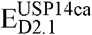and 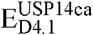, all E_D_-like states show two visible ubiquitin densities—one bound to RPN11 and conjugated to substrate and the other bound to USP14. In contrast, the S_D_-like states lack the densities of both RPN11-bound ubiquitin and substrates, as RPN11 is repositioned and blocks substrate entry into the OB ring. This difference implies that RPN11-catalysed deubiquitylation occurs exclusively along the substrate-engaged pathway and not in the substrate-inhibited pathway. In states E_A2.1_ and 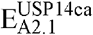, the Insert-1 (Ins-1) loop of RPN11 forms a β-hairpin sandwiched between the ubiquitin C-terminus and a segment of the rearranged N-loop from the RPT5 OB domain, forming a four-stranded β-sheet that was originally observed in state E_B_ without USP14 (Extended Data Fig. 7i)^9^. In the E_D_-like states, this β-sheet reduces to three β-strands as RPN11 is detached from the RPT5 N-loop by rotating toward the center of the OB ring (Extended Data Fig. 7h,i). The isopeptide bond linking ubiquitin to the substrate lysine remains uncleaved in these states (Extended Data Figs. 4b and 7h), indicating inhibited RPN11 activity and stalled substrate translocation. This arrangement is compatible with cryo-EM structures of the substrate-engaged yeast 26S proteasome inhibited by *o*PA^36^ (Extended Data Fig. 8a).

### Allosteric reprogramming of AAA-ATPase motor

The ten E_D_-like structures of the USP14^C114A^-proteasome visualize a continuum of conformational state transition around the AAA-ATPase ring, extending beyond previously reported states (Fig. 3). Conformational transitions from states 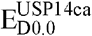 to 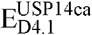 occur sequentially, as substrate detachment and ATP hydrolysis take place counterclockwise around the AAA-ATPase ring (Fig. 3d). In the newly observed major state 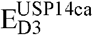, RPT3 is the only substrate-disengaged ATPase, and RPT4 is located at the top of the pore-1 loop staircase. Unlike previous studies where state 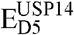 was identified, we did not observe it in this study despite exhaustive 3D classification, in which RPT2 or the RPT1-RPT2 dimer is expected to disengage from the substrate^29^. The interaction between the USP^C114A^ domain and the RPT1 OB domain remains relatively constant across states, while the distance from the USP^C114A^ domain to the RPT1 AAA domain varies significantly in the E_D_-like states but remains within a narrow range in the S_D_-like states (Fig. 2a,c). The widest configuration of the USP^C114A^-AAA interface is observed in state 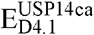, which closely resembles state 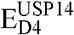 of the USP14^WT^-proteasome (Extended Data Fig. 8d). These differences in the conformational distribution suggest that USP14^C114A^ and inhibited RPN11 significantly reshapes the conformational landscape of the AAA-ATPase motor through allosteric regulation.

**Fig. 3:**
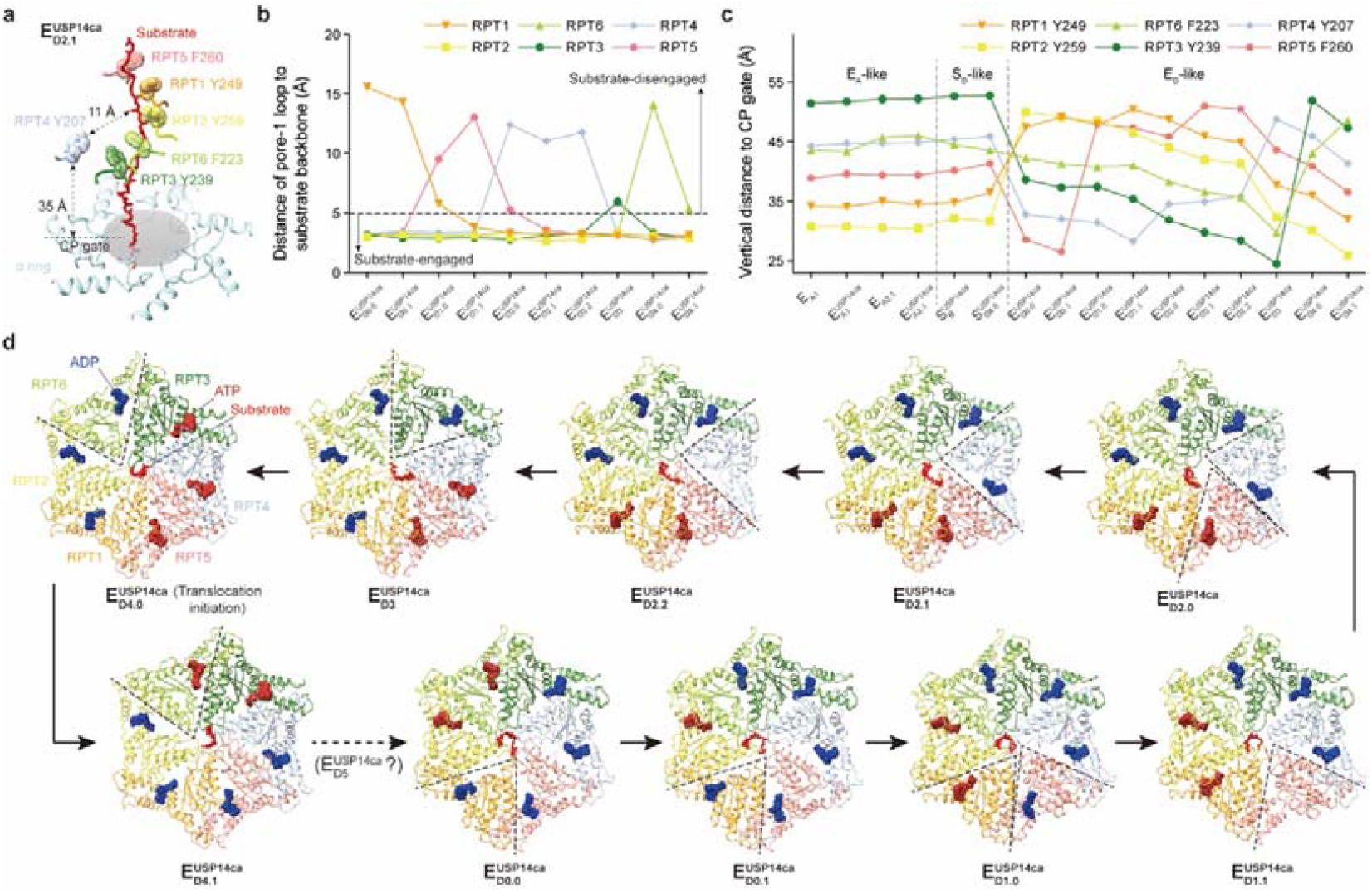
ATPase cycle in the USP14^C114A^-proteasome. **a**, T Typical architecture of the pore-1 loop staircase in state 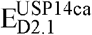. The substrate is modelled as a polypeptide backbone and is shown as stick representation. Side chains of the aromatic residues in the pore-1 loops are highlighted as transparent sphere representation. Segments of the CP α ring model are displayed, encircling the CP gate (grey ellipse). Distances from the RPT4 pore-1 loop to the substrate backbone and to the plane of the CP gate are indicated. **b**, Plot of the distances between the pore-1 loop of each ATPase and the substrate backbone across distinct states. Subunits with distances exceeding 5 Å (indicated by the dashed line) are considered substrate-disengaged. **c**, Plot of the distances between the pore-1 loop of each ATPase and the plane of the CP gate in distinct states. Only the aromatic residues of the pore-1 loops are considered in these distance measurements. **d**, Top views of atomic models of the AAA-ATPase ring in the ten E_D_-like states. Black arrows represent the deduced transitions between adjacent states, while the dashed arrow indicates a possible state 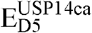 that is not observed in this study. The substrate-disengaged ATPases are marked by dashed angles. ATP (red) and ADP (blue) molecules in the nucleotide-binding pockets are highlighted using sphere representations.

The ATP-hydrolysis cycle within the AAA-ATPase motor is tightly coupled with substrate translocation in a highly coordinated process. The events of ATP binding, ADP release and ATP hydrolysis were reported to be allosterically synchronized in three adjacent ATPases^9,36^. However, our newly resolved E_D_-like structures reveal a different coupling pattern. ATP hydrolysis appears to occur in the ATPases separated by at least two subunits from those undergoing ADP release and ATP binding. Additionally, the entire AAA-ATPase ring harbors a higher proportion of ADP molecules compared to cryo-EM structures of the USP14^WT^-proteasome under native, non-disrupted states^9,29,36^ (Fig. 3d, Extended Data Fig. 6 and Table 2). For instance, state 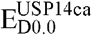 shares a similar overall conformation of the AAA-ATPase ring with state 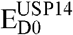 of the USP14^WT^ -proteasome^29^ (Extended Data Fig. 8f), but differs in the nucleotide-binding states of RPT1 and RPT3. While the conformation of the RPT3 AAA domain in state 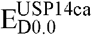 remains unchanged after ATP hydrolysis compared to state 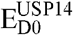 (Extended Data Fig. 8g,h), the dihedral angle in the RPT1 AAA domain is reduced by 10º due to delayed ADP release (Extended Data Fig. 8i,j).

In the wild-type proteasome, ADP release from the ATPase triggers a hinge-like motion between its small and large AAA subdomains, converting chemical energy from ATP hydrolysis into mechanical work that propagates around the AAA-ATPase ring for substrate unfolding and translocation^9^. However, in the RPN11-inhibited USP14^C114A^-proteasome, stalled substrate translocation disrupts the continuous flow of state transitions, halting allosteric communication between adjacent ATPases, delaying ADP release behind the schedule of ATPase cycle required for processive substrate degradation. As the AAA-ATPase motor attempts to overcome the stalled substrate translocation via continued ATP consumption in the USP14^C114A^-proteasome, more ADPs are accumulated in the AAA-ATPase ring (Fig. 3d).

### Pre-deubiquitylation bifurcation of substrate-processing pathways

The prominent differences in substrate densities between E_D_-like and S_D_-like states demonstrate that the catalytically deficient USP14^C114A^ directs the proteasomes along two distinct pathways— one leading to degradation of ubiquitylated substrates and the other toward rescuing and stabilizing deubiquitylated substrates. Systematic comparisons of the conformations reveal that while the AAA-ATPase ring remains consistent across the E_A_-like states, it undergoes notable variations in the E_D_-like states (Extended Data Fig. 7a-d). In the USP14^WT^-proteasome complex, state 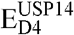 was previously interpreted as an intermediate bridging the E_A_-like and E_D_-like states during the initiation of substrate translocation^29^. Likewise, among the ten E_D_-like states of the USP14^C114A^-proteasome, state 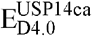 shows the closest resemblance to the E_A_-like states in its AAA-ATPase ring conformation, including the upturned RPT6 pore-1 loop (Extended Data Fig. 7a,e). On the other hand, state 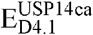 closely mirrors state 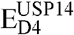 of the USP14^WT^-proteasome^29^ (Extended Data Fig. 8c). Therefore, state 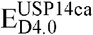 may represent a snapshot that precedes state 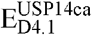 during the initiation of substrate translocation.

The conformational changes of the AAA-ATPase ring upon substrate insertion exhibit remarkable variations depending on the interaction with USP14. In the absence of USP14, the RPT6 AAA domain undergoes an outward rotation from states E_A2_ to E_B_, facilitating the opening of the central channel for substrate insertion (Extended Data Fig. 10a). This movement is concomitant with an outward shift in the RPT2 AAA domain, while the CP gate remains closed during state transition. By contrast, in the presence of USP14^C114A^, the RPT2 AAA domain experiences an upward shift from states 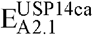 to 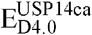 (Extended Data Fig. 10b), while the C-terminal tails of RPT1, RPT2, and RPT6 are inserted into the corresponding α-pockets of the CP, triggering the opening of the CP gate (Extended Data Fig. 5r).

### Allosteric control of substrate-processing pathways

To understand how USP14^C114A^ alters the state-transition pathway of the AAA-ATPase, we analyzed the inter-domain movements of individual ATPase subunits. In state 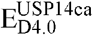, RPT1 undergoes ATP hydrolysis, resulting in an approximately 5° rotation between its large and small AAA subdomains compared to states 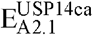 and E_B_ (Extended Data Fig. 10d). The RPT2 AAA domain exhibits only subtle changes among these states, characterized by a small displacement of approximately 3 Å in the short helix between Pro208 and Met214 (Extended Data Fig. 10c). This helix resides at the interface between RPT2 and the small AAA subdomain of RPT1, with the conformational change of RPT1 modulated by the USP^C114A^-RPT1 interaction. These observations imply that the USP^C114A^-RPT1 interaction induces a moderate conformational change within RPT1, which is then transmitted across the AAA-ATPase ring via inter-subunit interactions and amplified into a greater conformational change throughout the RP, ultimately leading to a distinct state-transition pathway for the entire complex.

The effect of USP14^C114A^-induced alteration of the state-transition pathway is also evident in RPT6. During the initiation of substrate translocation in the absence of USP14, the dihedral angle between the large and small AAA subdomains of RPT6 decreases after ADP release (E_A2_→E_B_ transition) and then increases upon ATP re-binding (E_B_→E_C1_ transition)^29^ (Extended Data Fig. 10e). In contrast, during the state transition of 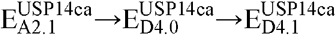, the small AAA subdomain of RPT6 rotates horizontally relative to the large AAA subdomain, in a direction orthogonal to that observed in the USP14-free E_A2_→E_B_→E_C1_ transition (Extended Data Fig. 10e). As a result, the dihedral angle between the large and small AAA subdomains of RPT6 remains almost unchanged, and the conformation of the RPT6 AAA domain in state 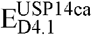 closely resembles that in state E_C1_, suggesting that the AAA-ATPase motor undergoes a smoother conformational transition to initiate substrate translocation in the presence of USP14^C114A^. This indicates that the proteasome bypasses the E_B_ and E_C_ states due to the allosteric regulation by USP14^C114A^, consistent with prior observation on the USP14^WT^-proteasome, which deviates from the RPN11-catalyzed state-transition pathway in the absence of USP14^29^.

### Fine-tuning the allocation of substrate-processing pathways

To explore the variability in how USP14 regulates proteasome dynamics, we conducted *in vitro* degradation assays and time-resolved cryo-EM analysis on the USP14^WT^-proteasome complex using the model substrate Ub_*n*_-Sic1^PY^. Notably, we observed faster substrate degradation in a buffer containing 100 mM NaCl compared to a NaCl-free buffer, indicating reduced suppression of proteasome activity by USP14 (Fig. 4a). Using time-resolved cryo-EM data, we reconstructed the conformational distributions of the proteasome during substrate degradation at two different NaCl concentrations (Fig. 4b and Extended Data Fig. 11). For both conditions, the population of the E_D_-like states increased as the reaction progressed, peaking at approximately 1 minute, and then decreased due to substrate depletion. However, in the presence of 100 mM NaCl, the population of the E_D_-like states reached a higher peak, while the population of the S_D_-like states remained consistently lower, indicating that the USP14^WT^-proteasome under this condition preferred the substrate-engaged pathway over the substrate-inhibited pathway.

**Fig. 4:**
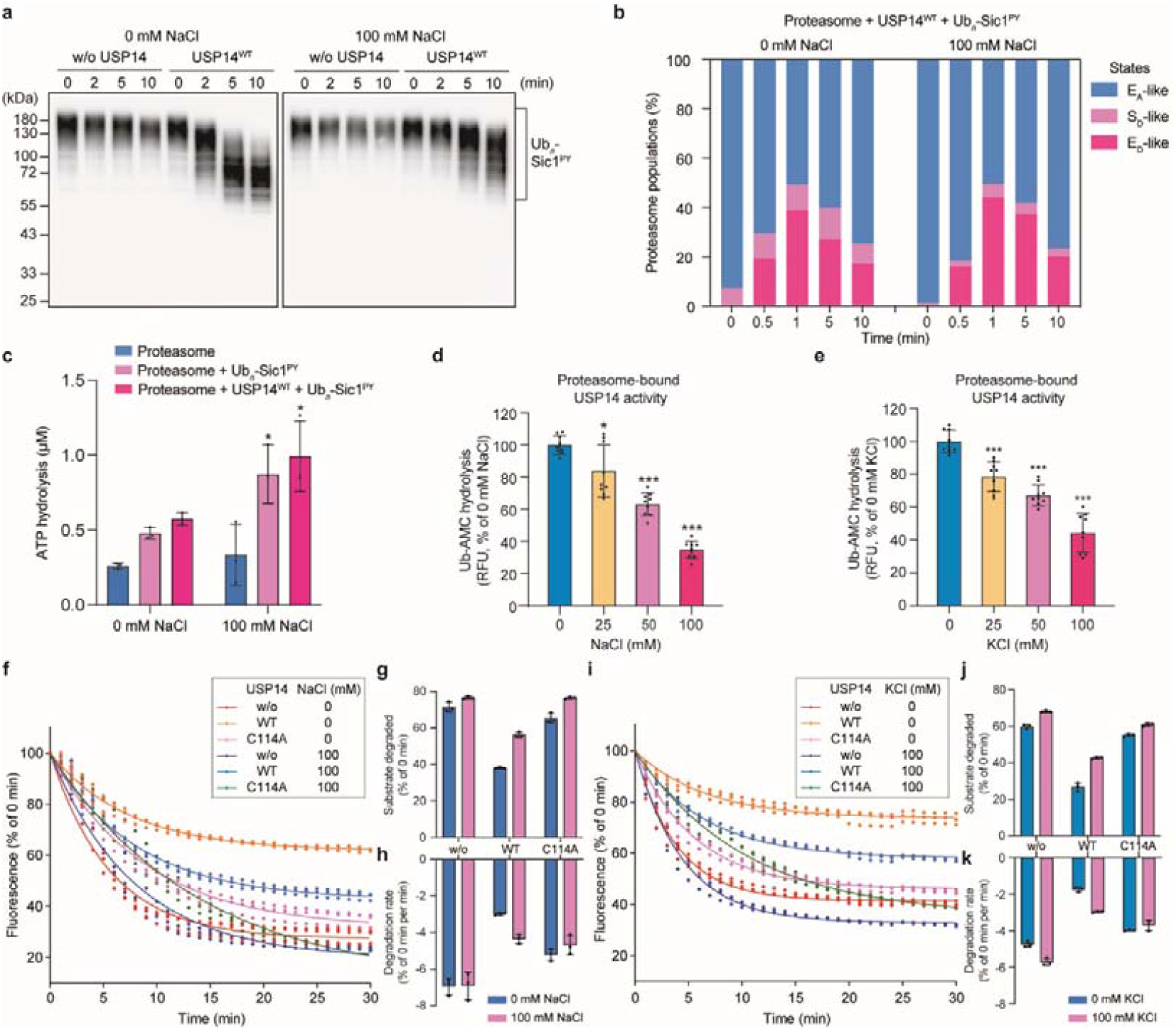
Salt-dependent modulation of substrate processing by the USP14-proteasome complex. **a**, *In vitro* degradation of T7-tagged Ub_*n*_-Sic1^PY^ by the human proteasome at 10 □, analyzed by SDS-PAGE and western blot with anti-T7 antibody. **b**, Kinetic changes in the overall populations of proteasome particles in E_A_-like, S_D_-like and E_D_-like states, as determined by time-resolved cryo-EM analysis of the proteasome samples at 10 □. Control data under the NaCl-free condition were sourced from our previous study^29^. The particle number data used in this histogram are provided in Extended Data Fig. 11b,c. **c**, ATPase activity of the proteasome in the presence or absence of USP14 or substrate at different NaCl concentrations. Data are shown as mean ± s.d. from three independent experiments, with each experiment including three replicates. \**p*<0.05 compared to 0 mM NaCl, determined using a two-tailed unpaired *t*-test. **d, e**, The DUB activity of USP14 in the presence of the proteasome at different NaCl (**d**) or KCl (**e**) concentrations. RFU, relative fluorescence units. The data are presented as mean ± s.d. from three independent experiments, each consisting of three replicates. \**p*<0.05, \*\*\**p*<0.001 compared with 0 mM NaCl using a two-tailed unpaired *t*-test. **f**-**k**, *In vitro* degradation of the Ub_*n*_-NtSic1^PY^-cp8sGFP substrate by the proteasome in the presence or absence of USP14 variants at different NaCl (**f**) or KCl (**i**) concentrations at 30 □. The dots indicate percentages of initial fluorescence for each condition, derived from three independent replicates, and the decay curves are global fits to single exponentials. This assay was validated using western blot (Extended Data Fig. 1d). The total percentage of degraded substrates and the rate of degradation in the first 10 mins are compared side-by-side in separate plots in panels **g** and **h** for different NaCl concentrations, respectively, and in panels **j** and **k** for different KCl concentrations, respectively.

To visualize the dynamic conformational space of the proteasome, we applied deep learning-based manifold embedding to analyze the time-resolved cryo-EM data, mapping single-particle images onto a series of two-dimensional landscapes representing the proteasome’s conformational dynamics over time (Methods and Extended Data Fig. 12a-c). This approach allowed us to compare the dynamic evolution of conformational landscape under different buffer conditions (Fig. 5 and Extended Data Fig. 12d-g). In the NaCl-free condition, the S_D_-like states occupied a larger, lower energy region of the landscape, indicating that these states were up-regulated and more stable. In contrast, the presence of 100 mM NaCl resulted in a more enduring population of the E_D_-like states, while the S_D_-like states were barely detectable and apparently down-regulated, consistent with reduced suppression of the proteasome activity observed in the degradation assays.

**Fig. 5:**
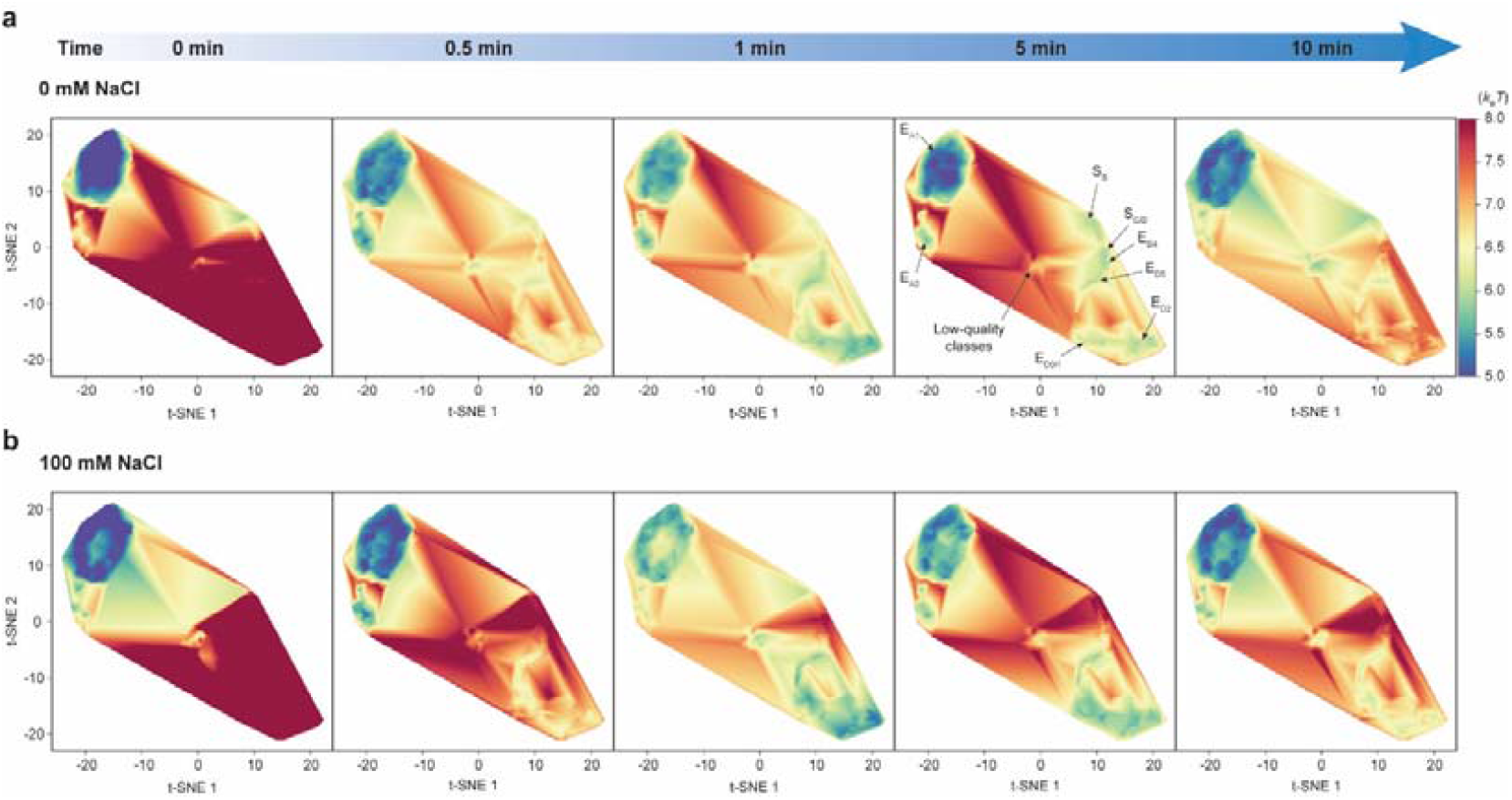
Salt-dependent variation of time-resolved energy landscapes of the USP14-proteasome complex during substrate degradation. **a, b**, Energy landscapes at different time points, reconstructed from time-resolved cryo-EM experiments in the absence of NaCl (**a**) and in the presence of 100 mM NaCl (**b**). The same t-SNE coordinates and color bar are used for plotting the energy landscapes in all panels. The dataset for panel (**a**) is from our previous study^29^.

### Opposing effects of salt-dependent modulation on USP14 and the proteasome

Intriguingly, the proteasome exhibits higher rates of both Ub_*n*_-Sic1^PY^ substrate degradation and ATP hydrolysis under 100 mM NaCl, regardless of whether USP14 was present (Fig. 4a,c). Ubiquitin constantly uses its positively charged surface surrounding residue Ile44 to interact with the negatively charged ubiquitin-binding sites on the ubiquitin receptors RPN1, RPN10 and RPN13, as well as RPN11^16,24,29,39^. Because electrostatic interactions contribute to ubiquitin recognition by the proteasome, sodium ions are expected to modulate ubiquitylated substrate interactions with the proteasome via the Debye screening effect. Moreover, different ionic strengths can also impact the inter-subunit interactions within the proteasome, as well as nucleotide binding and release in the AAA-ATPase motor.

To determine whether and how the proteasome’s sensitivity to ionic strength is related to USP14, we further analyzed the DUB activity of USP14 under varying salt concentrations. Interestingly, Ub-AMC hydrolysis assays revealed that the basal DUB activity of free USP14 remains largely unaffected by NaCl or KCl (Extended Data Fig. 1f,g). However, when USP14 was bound to the proteasome, its DUB activity was negatively correlated with NaCl or KCl concentration (Fig. 4d,e). Moreover, microscale thermophoresis (MST) measurements showed no apparent decrease in the binding affinity of USP14 to the proteasome in the presence of 100 mM NaCl (Extended Data Fig. 1e), suggesting that the population of USP14-bound proteasome was unlikely to diminish under these conditions with moderate ionic strength. Thus, we reason that salt-enhanced proteasome activity allosterically down-regulates the DUB activity of USP14, while the reduced DUB activity of USP14 conversely contributes to reduced proteasome suppression. Both effects favor the substrate-engaged pathway over the substrate-inhibited pathway, giving rise to enhanced degradation performance.

Finally, we confirmed these findings using real-time quantitative fluorescence monitoring of proteasomal degradation of another model substrate, Ub_*n*_-NtSic1^PY^-cp8sGFP, which was used in cryo-EM analysis (Fig. 4f,i). The fluorescence decay rate and final fluorescence levels indicated the degradation rate and the amount of surviving or partially degraded substrates, respectively. Under 100 mM NaCl or KCl, the USP14^WT^-bound proteasome exhibits a faster degradation rate and fewer remaining substrates compared to the NaCl/KCl-free condition (Fig. 4g,h,j,k). This result aligned well with the degradation assays of Ub_*n*_-Sic1^PY^, indicating that proteasome suppression by USP14 was reduced when proteasome activity was enhanced by NaCl or KCl. Moreover, the USP14^C114A^-bound proteasome degraded substrates more slowly than the USP14-free proteasome, reflecting non-catalytic suppression of proteasome activity by USP14. The addition of NaCl or KCl results in more substrates degraded by the USP14^C114A^-bound proteasome, further demonstrating that USP14’s non-catalytic regulation of proteasome activity can be fine-tuned by ionic strength of sodium or potassium. Taken together, our findings reveal that the non-catalytic regulation of USP14 on proteasome activity can be modulated by changes in ionic strength of sodium or potassium, providing insight into how proteasome dynamics are fine-tuned under different physiological conditions.

## Discussion

Both sodium and potassium are physiologically important electrolytes to maintain the resting membrane potential and transmembrane electrochemical gradient necessary for cell signaling and regulation of membrane protein functions. In neurons, sodium and potassium are vital for generating and propagating action potentials, whereas in muscle cells, they are critical for regulating excitation-contraction coupling, which converts a neural signal into muscle contractions. Therefore, understanding how sodium and potassium impact the proteasome function is physiologically significant. High salt concentrations can induce protein dissociation from complex assemblies, such as the dissociation of USP14 from the proteasome^32^. Therefore, our previous cryo-EM studies of USP14-bound proteasome structures were performed in the absence of NaCl or KCl to maximize holoenzyme stability^29^. In contrast, all cryo-EM structures of substrate-engaged USP14-proteasome in this study were determined in the presence of 100 mM NaCl in the buffer. We systematically analyzed the dependence of the 26S proteasome on sodium and potassium concentrations in the range that closely mimics *in vivo* conditions. The findings that the proteasome activity is upregulated, while USP14 activity is downregulated, with increasing sodium and potassium concentrations, respectively, have important physiological implications for understanding the proteostasis and proteasome regulation in dynamic intracellular environment. The substrate-processing pathways regulated by USP14 in the proteasome operate largely independently of its catalytic DUB activity but are notably modulated by salt concentration. USP14 allosterically influences the state transitions of the AAA-ATPase motor, thereby regulating the allocation between substrate-inhibited and substrate-engaged pathways within the proteasome population. By adjusting sodium and potassium concentrations, the probabilistic partitioning between these two opposing pathways can be fine-tuned, revealing a previously unappreciated regulatory mechanism and adding another layer of complexity to proteasome biology. This raises the question of whether other metal ions, such as calcium, may also contribute to proteasome regulation, warranting further mechanistic investigation.

In contrast to prior studies that only captured the post-deubiquitylation snapshots^29^, our cryo-EM structures of the RPN11-inhibited USP14^C114A^-proteasome complex provide a systematic view of conformational states just before the DUB-catalyzed deubiquitylation. The C114A mutation in USP14, along with RPN11 inhibition, induces certain structural features of the proteasome that deviate from physiologically native conditions, potentially reflecting off-pathway effects, such as enriched ADP content or delayed nucleotide exchange in the ATPases. Despite these deviations, the overall substrate-engaged AAA-ATPase conformations in all E_D_-like states observed here align well with the asymmetric hand-over-hand model of substrate translocation^17,18^ and can be interpreted as intermediate conformations along the functional pathway of state transitions (Figs. 3d and 6a). These newly identified conformational states fill in the major missing gaps in previously published snapshots of substrate-engaged human proteasome^9,29,36^, completing the characterization of all major intermediate states around the ATPase ring and further corroborating the asymmetric hand-over-hand mechanism for substrate translocation (Fig. 6a,b).

**Fig. 6:**
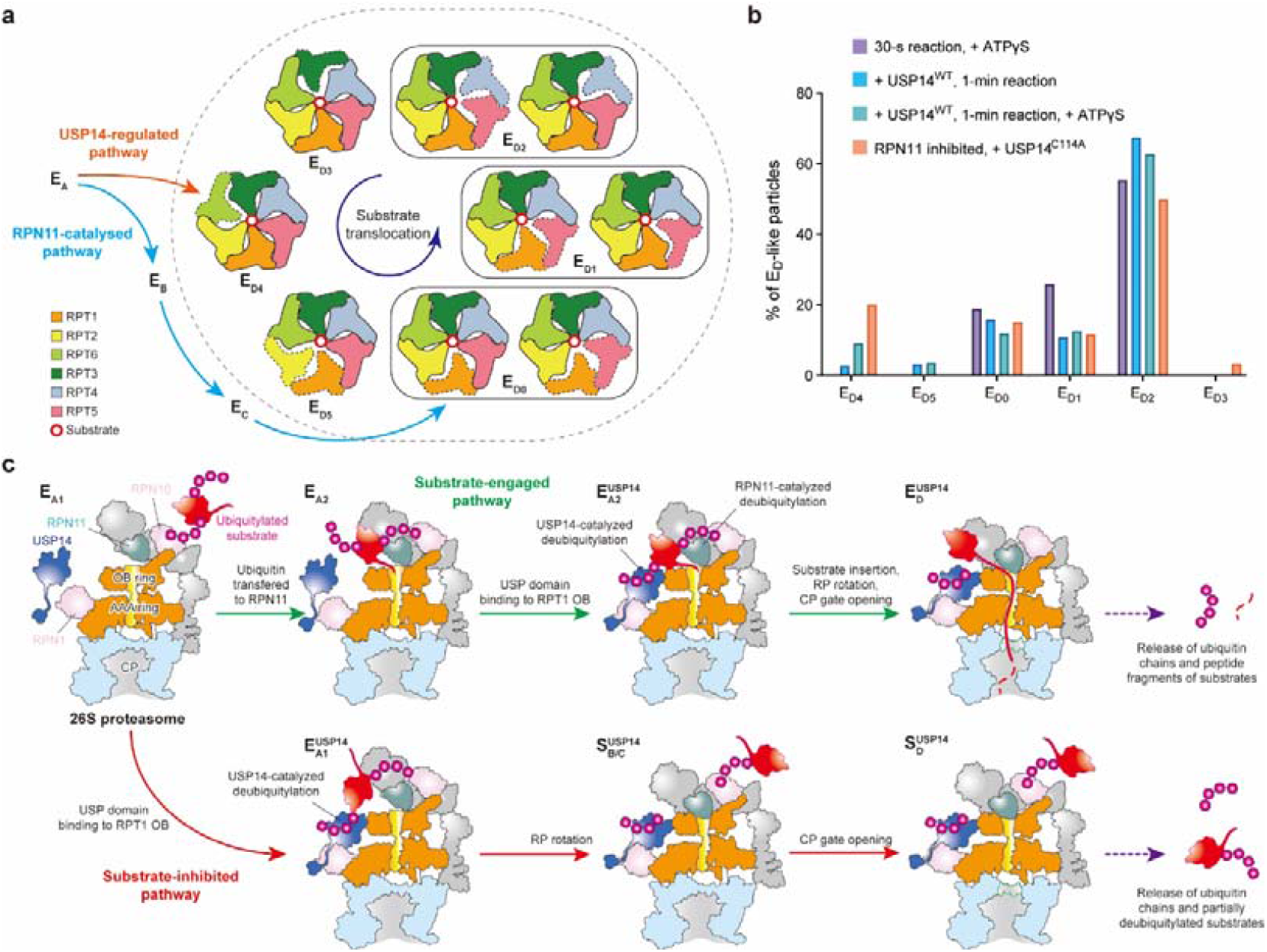
Model for state transitions and pathway selection by the USP14-proteasome complex. **a**, The illustration depicts the complete transition cycle through all major conformational states of the substrate-engaged AAA-ATPase ring during processive substrate translocation, based on cryo-EM studies of both USP14-free^9,40^ and USP14-bound proteasomes (ref.^29^ and this study). The dashed outline encompasses a series of sub-E_D_ states characterized by a substrate strand within the central channel and an open CP gate. Substrate-disengaged ATPases are marked with dashed edges. In the absence of USP14, the proteasome follows a substrate degradation pathway where deubiquitylation is catalyzed by RPN11 (blue arrows). The proteasome transitions from state E_C_ into state E_D0_, opening the CP gate and entering the ATPase cycle for processive substrate translocation and degradation. In contrast, in the presence of USP14 (orange arrow), the proteasome bypasses states E_B_ and E_C_, directly entering the ATPase cycle for substrate translocation in state E_D4_. **b**, The distribution of each major substrate-engaged state across the total number of E_D_-like particles under different conditions, as determined by previous cryo-EM studies^29,40^ and this study. **c**, Proposed model for pathway selection and allocation by the proteasome in the presence of USP14. The proteasome can transition into either the substrate-engaged pathway (green arrows) or the substrate-inhibited pathway (red arrows), depending on the allosteric competition between USP14 and the proteasome. As indicated by the dashed arrows on the right, these pathways result in opposing outcomes for the processed substrates being either degraded or rescued.

Our data clarify that the non-catalytic, allosteric regulation by USP14 alone can induce both substrate-inhibited and substrate-engaged pathways in parallel (Fig. 6c). The initial insertion of a substrate into the AAA-ATPase motor competes thermodynamically and kinetically with the recognition of substrate-conjugated ubiquitin by USP14. If USP14 recognizes ubiquitin before substrate engagement with the ATPases, the substrate-inhibited pathway is activated, allowing USP14 to stabilize the substrate through deubiquitylation. Conversely, faster substrate engagement triggers the pathway for substrate degradation, regardless of the presence of USP14. Thus, USP14 can non-catalytically promote the substrate-inhibited pathway rather than directly abolishing RPN11 activity^33,34^. On the substrate-engaged pathways, USP14 allosterically modulates the inherently imbalanced statistical distribution of proteasome states through multi-interfacial interactions^29^ (Fig. 6b). The initiation of substrate translocation is allosterically reprogrammed to occur during the E_A2_-to-E_D4_ transition, whereas without USP14 it takes place during the E_C_-to-E_D0_ transition^9,40^ (Fig. 6a). Due to its unique spatial arrangement in the proteasome, USP14 reorients the substrate to an alternative insertion path, preventing RPN11 from capturing ubiquitin on the same substrate. Consequently, USP14 is poised to deubiquitylate additional substrates aside from the one being actively degraded by the proteasome, potentially reducing the pool of degradable substrates and suppressing overall degradation. These non-catalytic properties are indispensable for timing the catalytic function of USP14 on the proteasome, positioning it as a highly efficient proofreading or “error correction” gatekeeper that antagonizes RPN11 to fine-tuning substrate processing.

## Methods

### Protein Expression and purification

Human 26S proteasome was affinity-purified from a stable HEK293 cell line with HTBH (6×His, TEV cleavage site, biotin and 6×His)-tagged RPN11 (a gift from L. Huang) as previously described^8,9,11,29^. Briefly, harvested cells were lysed by a Dounce homogenizer in lysis buffer (50 mM PBS [77.4% Na_2_HPO_4_, 22.6% NaH_2_PO_4_, pH 7.4], 5 mM MgCl_2_, 5 mM ATP, 0.5% NP-40, 1 mM DTT and 10% glycerol) containing a protease inhibitor cocktail. After centrifugation, the supernatant was incubated with Streptavidin Agarose Resin 6FF (Yeasen) for 3 hours at 4 °C. The resin beads were washed with 20 bed volumes of lysis buffer supplemented with 100 mM NaCl to remove endogenous USP14 associated with the proteasome. The proteasome was subsequently cleaved from the beads using TEV protease (Invitrogen) and further purified via size-exclusion chromatography using a Superose 6 10/300 GL column (GE Healthcare) pre-equilibrated with FPLC buffer (30 mM HEPES [pH 7.5], 60 mM NaCl, 0.5 mM DTT, 1 mM MgCl_2_, 0.6 mM ATP, 10% glycerol).

In this paper, we used two types of model substrates, both fused to 6×His and T7 tags: Sic1^PY^ and NtSic1^PY^-cp8sGFP. The PY motif (Pro-Pro-Pro-Ser) inserted at the N-terminus of these substrates facilitates recruitment of the E3 ubiquitin ligase WW-HECT (derived from Rsp5 of *Saccharomyces cerevisiae* through the deletion of its N-terminal 220 amino acids) to catalyze polyubiquitylation^41^. The recombinant substrate NtSic1^PY^-cp8sGFP consists of an N-terminal Sic1^PY^-derived unstructured region (68 amino acids) and a C-terminal cp8sGFP, a circular permutant of superfold GFP in which the 8th β-strand serves as the C-terminus instead of the 11th. This variant is well-folded but less stable than superfold GFP^42,43^) with all cysteines substituted with alanines to prevent the formation of disulfide bonds. This newly designed substrate has a similar sequence length (316 amino acids) and molecular weight (34.5 kDa) to Sic1^PY^, and contains multiple lysines as ubiquitylation sites (Extended Data Fig. 1a,c).

Wild-type USP14 (USP14^WT^), USP14^C114A^ (an inactive variant with the catalytic cysteine substituted with alanine), UBE1, UBCH5A, WW-HECT and Sic1^PY^ were recombinantly expressed in BL21-CondonPlus (DE3)-RIPL cells (Shanghai Weidi) and purified as previously described^29^. For the purification of NtSic1^PY^-cp8sGFP, the plasmid (pET-21a-NtSic1^PY^-cp8sGFP, cloned by GenScript) was transformed into BL21-CondonPlus (DE3)-RIPL cells (Shanghai Weidi). The cells were grown to an OD600 of 0.6-0.7 in LB medium and induced by 0.2 mM IPTG overnight at 20 °C. The cells were then harvested, resuspended in lysis buffer (25 mM Tris-HCl [pH 8.0], 150 mM NaCl, 0.2% Triton-X-100, 1 mM TCEP, 10% glycerol and protease inhibitor cocktail) and lysed by sonication. The supernatant was incubated with pre-equilibrated HisSep Ni-NTA Agarose Resin 6FF (Yeasen) for 2 hours at 4 °C. The resin beads were washed with 20 bed volumes of washing buffer (50 mM Tris-HCl [pH 8.0], 150 mM NaCl, 10% glycerol, 1 mM TCEP and 20 mM imidazole), followed by elution with buffer containing 150 mM imidazole and the same other ingredients as the washing buffer. The eluted proteins were further purified by size-exclusion chromatography using a Superdex 75, 10/300 GL column (GE Healthcare) pre-equilibrated with FPLC buffer (50 mM Tris-HCl [pH 8.0], 150 mM NaCl, 1 mM DTT and 10% glycerol).

### Preparation of polyubiquitinated substrates

Ubiquitylation of Sic1^PY^ was performed as previously described^29^. For the ubiquitylation of NtSic1^PY^-cp8sGFP, a mixture containing 1.2□μM NtSic1^PY^-cp8sGFP, 0.5□μM UBE1, 2□μM UBCH5A, 1.4□μM WW-HECT and 1□mg mL^−1^ ubiquitin (Boston Biochem) was incubated in reaction buffer (50□mM Tris-HCl [pH 7.5], 100□mM NaCl, 10□mM MgCl_2_, 2□mM ATP, 1□mM DTT and 10% glycerol) for 3□hours at room temperature. The polyubiquitylated NtSic1^PY^-cp8sGFP (Ub_*n*_-NtSic1^PY^-cp8sGFP) was purified by incubating with Ni-NTA Agarose Resin 6FF (Yeasen) at 4□°C for 1□hour. The resin was washed with 20 bed volumes of washing buffer (50□mM Tris-HCl [pH 7.5], 100□mM NaCl, 10% glycerol and 15 mM imidazole). Ub_*n*_-NtSic1^PY^-cp8sGFP was then eluted using elution buffer (150 mM imidazole in the same buffer as the washing buffer), concentrated and exchanged to the storage buffer (50□mM Tris-HCl [pH 7.5], 100□mM NaCl, 1 mM TCEP and 10% glycerol) using Zeba™ Spin Desalting Columns with a 7K molecular weight cut-off (Invitrogen).

### Liquid chromatography-tandem mass spectrometry

Liquid chromatography-tandem mass spectrometry (LC-MS/MS) assay was performed to analyze the ubiquitylation of NtSic1^PY^-cp8sGFP. Briefly, ubiquitylated NtSic1^PY^-cp8sGFP proteins were treated with 5 mM DTT and 10 mM iodoacetamide (IAA), then digested into peptides using trypsin. The enzymatic reaction was terminated by 0.1% trifluoroacetic acid (TFA). LC-MS/MS analysis was conducted using an Easy-nLC 1200 system (Thermo Fisher), a C18 column (Acclaim PepMap 100, 75 μm × 2 cm nanoViper, Thermo Fisher) and an Orbitrap Fusion Lumos (Thermo Fisher). Peptides were loaded onto the C18 column (Thermo Fisher) at a speed of 280 nL/min and separated using a segmented gradient. Solvent A contains 0.1% (v/v) formic acid in H_2_O and solvent B consisted of 0.1% (v/v) formic acid and 80% acetonitrile. The gradient was set as follows: 4%–8% solvent B in 5 minutes; 8% 20% solvent B in 45 minutes; 20%–30% solvent B in 10 minutes; 30%–90% solvent B in 13 minutes; 90% solvent B in 3 minutes. The mass spectrometer was operated in data-dependent mode. MS1 scans were conducted using an Orbitrap Mass Analyzer across a range of 300-1500 m/z with a resolution of 120K at m/z 200, an automatic gain control (AGC) target at 5 × 10^5^, and a maximum injection time at 90 ms, followed by MS2 scans generated by HCD (30% energy) fragmentation at the resolution of 30K, AGC target 5 × 10^4^, and a maximum injection time of 100 ms. The generated data were processed using Proteome Discoverer v.2.2 software (Thermo Fisher). Carbamidomethylation (at Cys) was set as fixed modification and ubiquitylation (at Lys) was set as variable modification.

### In vitro degradation assay

Human proteasomes (∼30 nM) were incubated with Ub_*n*_-Sic1^PY^ or Ub_*n*_-NtSic1^PY^-cp8sGFP (∼300 nM) in degradation buffer (50 mM Tris-HCl [pH 7.5], 5 mM MgCl_2_ and 1 mM ATP; supplemented with 100 mM NaCl or 100 mM KCl in some cases). Purified recombinant USP14^WT^ or USP14^C114A^ (∼1.2 μM) were incubated with the proteasome for 15 minutes before initiating the degradation. For the degradation of Ub_*n*_-Sic1^PY^, the reaction mixtures were incubated at 10 °C for 0, 2, 5 and 10 minutes, then terminated by the addition of SDS loading buffer and analyzed via western blot using an anti-T7 antibody (Beyotime Biotechnology, 1:1000 dilution). For the degradation of Ub_*n*_-NtSic1^PY^-cp8sGFP, the reaction mixtures were incubated at 30 °C for 0, 5, 15 and 30 minutes, and then terminated by the addition of SDS loading buffer and analyzed via western blot using an anti-T7 antibody. Alternatively, the degradation assay was conducted through fluorescence monitoring. Proteasomes were incubated with USP14 in degradation buffer containing 1 mg mL^-1^ BSA for 15 min at room temperature. Ub_*n*_-NtSic1^PY^-cp8sGFP was then added to the reaction mixtures, and the fluorescence of cp8sGFP (488 nm excitation; 520 nm emission) was measured every minute over a 30-minute period using the Varioskan Flash spectral scanning multimode reader (Thermo Fisher).

### Microscale thermophoresis (MST)

The human proteasomes were labelled using the Monolith NT™ Protein Labeling Kit (NanoTemper) and exchanged to reaction buffer (50 mM Tris-HCl [pH 7.5], 100 mM NaCl, 5 mM MgCl_2_ and 1 mM ATP) supplemented with 0.05% Tween-20. To study the interaction between the proteasome and USP14, a concentration series of USP14 was prepared using a 1:1 serial dilution in the reaction buffer. The interaction was initiated by adding an equal volume of the labelled proteasome to each serial dilution of USP14, and the measurement was taken using the MonolithTM NT.115 (NanoTemper) at 20% LED excitation power and 40% MST power.

### Ubiquitin-AMC hydrolysis assay

A ubiquitin-AMC (Ub-AMC; Boston Biochem) hydrolysis assay was performed to verify the deubiquitylation activity of USP14 in the presence of different concentrations of NaCl or KCl. 1 nM Ub-VS-treated proteasome and 0.2 μM USP14 were incubated in reaction buffer (50 mM Tris-HCl [pH 7.5], 5 mM MgCl_2_, 1 mM ATP, 1 mM DTT, 1 mM EDTA and 1 mg mL^-1^ ovalbumin [Diamond]) supplemented with 0 mM, 25 mM, 50 mM or 100 mM NaCl or KCl for 15 minutes at room temperature. The reaction was initiated by adding 1 μM Ub-AMC (BioVision) to the mixture and incubated for 30 minutes at 30 °C. Fluorescence was measured using the Varioskan Flash spectral scanning multimode reader (Thermo Fisher). For the deubiquitylation activity of free USP14, the reaction was conducted using 1 μM USP14 incubated with 0.5 μM Ub-AMC in the absence of the proteasome.

### ATPase activity assay

The ATPase activity of the proteasome was determined using malachite green phosphate assay kits (Sigma). 30 nM proteasome and 1.2 μM USP14 were incubated in reaction buffer (50 mM Tris-HCl [pH 7.5], 100 mM NaCl, 5 mM MgCl_2_ and 0.5 mM ATP) for 15 minutes. 300 nM Ub_*n*_-Sic1^PY^ was added to the mixture and incubated for 5 minutes at 10°C. Then malachite green buffer was added and the mixture was incubated for an additional 30 minutes at room temperature. The absorbance at 620 nm (OD620) was recorded using a Varioskan Flash spectral scanning multimode reader (Thermo Fisher).

### Sample and grid preparation for cryo-EM

In this study, we prepared cryo-EM grids for two types of samples: the time-resolved sample and the USP14^C114A^-proteasome sample. For the time-resolved sample, we performed cryo-EM sample preparation as previously described^29^, except for the NaCl concentration. All purified proteins were exchanged into the same imaging buffer (50□mM Tris-HCl [pH 7.5], 100 mM NaCl, 5□mM MgCl_2_ and 1□mM ATP). A mixture of 1 μM human proteasomes and 10□μM USP14 was incubated for 20□minutes at 30□°C, then cooled to 10□°C. The mixture was divided into 5 PCR tubes, and 10□μM Ub_*n*_-Sic1^PY^ was added to four of these tubes and incubated at 10□°C for 0.5, 1, 5 and 10□minutes, respectively. For the control and initial state (0 minutes) of the degradation reaction, the sample in the last tube was diluted with the imaging buffer to match the concentrations of the proteasome in other tubes. After incubation, 4.5 μl of each sample was applied to glow-discharged Quantifoil R1.2/1.3 holey carbon grids. NP-40 was added to the reaction mixture to a final concentration of 0.005% immediately before cryo-plunging to prevent orientation preference. These samples were used for comparison with the previous time-resolved cryo-EM study, in which Ub_*n*_-Sic1^PY^ was degraded by the USP14-proteasome complex in a NaCl-free reaction buffer^29^.

For the USP14^C114A^-proteasome sample, we exchanged all purified proteins to the imaging buffer (50□mM Tris-HCl [pH 7.5], 100 mM NaCl, 5□mM MgCl_2_ and 1□mM ATP) containing the RPN11 inhibitor ortho-phenanthroline (*o*PA) at 5 mM. A mixture of 1 μM of the human proteasome and 10□μM of the inactive USP14^C114A^ variant was incubated for 20□minutes at room temperature. Afterward, 10□μM of Ub_*n*_-NtSic1^PY^-cp8sGFP was added to the mixture and incubated for 10 minutes at 10□°C, and NP-40 was added to a concentration of 0.005%. A total of 4.5 μl of the final sample was applied to a glow-discharged Quantifoil R1.2/1.3 holey carbon grid for cryo-plunging. In addition to RPN11 and USP14, the purified human proteasome may be associated with another DUB, UCH37 (also known as UCHL5), which was demonstrated to exclusively cleave branched ubiquitin chains bearing K48 linkages^44^. Since the E3 ligase WW-HECT preferentially catalyzes Lys63-linked ubiquitylation^45^, the substrate used in this study can hardly be targeted by UCH37. Therefore, the entire DUB activity of the USP14^C114A^-proteasome complex can be considered inactive.

### Cryo-EM data collection

Cryo-EM data were collected on the Titan Krios G2 microscope (Thermo Fisher), operated at 300 kV and equipped with a post-column BioQuantum energy filter (Gatan) and a K2 Summit direct electron detector (Gatan). Automated data acquisition was performed using SerialEM^46^ in super-resolution counting mode, with an energy slit width of 20□eV and a nominal defocus set between -1.0 and -2.0□μm. Each micrograph was recorded as a 40-frame dose-fractioned movie stack, with a total exposure time of 10 seconds and a cumulative electron dose of ∼50 electrons per Å^2^. For the time-resolved sample cryo-plunged at reaction times of 0, 0.5, 1, 5 and 10□minutes, the numbers of collected micrographs were 1,397, 2,438, 4,913, 2,224 and 2,280, respectively, with calibrated super-resolution pixel sizes of 0.685 Å. For the USP14^C114A^-proteasome sample, a total of 17,040 micrographs was collected at a calibrated super-resolution pixel size of 0.668 Å.

### Cryo-EM data processing

Dose-weighted motion correction was performed using MotionCor2^47^ and contrast transfer function (CTF) estimation was done by Gctf^48^. Particles were automatically picked using DeepEM^49^ and manually checked in EMAN2^50^. Reference-free 2D and 3D classification, focused 3D classification, aberration correction, per-particle CTF refinement and high-resolution auto-refinement were carried out on two-fold binned particles (with pixel sizes of 1.37 Å for the time-resolved dataset and 1.336 Å for the USP14^C114A^-proteasome dataset) using RELION 4.0^51^. Particle subtraction and re-centering were performed using custom Python scripts invoking RELION 4.0. Conformational changes were analyzed through in-depth 3D classification using AlphaCryo4D^52^.

Given the highly complex conformational heterogeneity of the proteasome, a hierarchical 3D classification strategy was applied to analyze the datasets as previously described^29^ (Extended Data Fig. 2). First, doubly capped proteasome particles (RP-CP-RP) were separated from singly capped ones (RP-CP) to generate pseudo-singly capped (PSC) particles by re-centering the particle box. Second, these PSC particles were subjected to CP-masked auto-refinement followed by aberration and CTF refinement. Then, alignment-skipped RP-focused 3D classification was performed to separate E_A_-like particles from E_D_/S_D_-like ones (E_A_, E_D_ and S_D_ denote ubiquitin-recognition state, substrate-engaged state and substrate-inhibited state, respectively). Particles in E_D_/S_D_-like states can be easily distinguished from those in E_A_-like states due to the dramatic rotation of the RP. Third, CP-masked auto-refinement, followed by RP-focused 3D classification and conformational landscape analysis by AlphaCryo4D, was performed to further classify the RP conformations into several clusters. Fourth, subtraction of the CP density and box re-centering to the RP were performed for PSC particles, generating CP-subtracted particles. These particles were then subjected to several rounds of alignment-skipped RP-focused 3D classification, resulting in tens of different conformational states with distinguishable structural features in the overall RP, the AAA-ATPase ring and the substrate. Final high-resolution reconstructions of these states were performed on PSC particles by refining RP and CP independently and merging the two focused maps in Phenix^53^.

For the time-resolved dataset, the aim was to compare results with prior studies rather than data processing at achieving high-resolution reconstructions but rather for comparison with previous studies. Data collected at different time points were combined and processed together until classified into distinct conformational states (Extended Data Fig. 11a). Then, these data were separated according to their time labels to calculate conformational distributions for each time point. A total of 309,819 E_A_-like particles and 154,863 E_D_/S_D_-like particles were obtained across all time points after step 2. Surprisingly, the proportion of proteasome particles in the substrate-inhibited S_D_-like state was much lower compared to our previous study in which the same substrate was degraded by the USP14-bound proteasome in a NaCl-free buffer^29^. Since the conformations of the E_D_-like maps closely resembled those of the S_D_-like maps, and the total number of E_D_-like and S_D_-like particles was insufficient to ensure the classification accuracy, we used a seed-guided 3D classification method for further analysis and verification (Extended Data Fig. 11a). Particles from this dataset were combined with those from the previous study^29^, using the latter as seeds to attract particles with similar conformational characteristics through repeated focused 3D classifications. There were four intermediate S_D_-like states reported in the previous study^29^, from which 43,100 particles in state 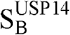, 21,071 particles in state 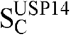, 37,985 particles in state 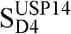 and 33,590 particles in state 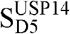 were selected and combined with E_D_/S_D_-like particles in this study. Each combination was subjected to focused 3D classification five times, after which particles assigned to classes corresponding to the states of the seed particles at least three times were classified as S_D_-like particles. Finally, a total of 18,239 S_D_-like particles and 136,624 E_D_-like particles were identified in the time-resolved dataset in this study, indicating that the S_D_-like states became less populated in the presence of 100 mM NaCl in the buffer.

For the USP14^C114A^-proteasome dataset, the hierarchical 3D classification strategy yielded 16 conformational states of the RPN11-inhibited USP14^C114A^-proteasome complex (Extended Data Fig. 2), including four E_A_-like states (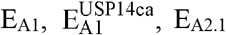 and 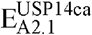), two S_D_-like states (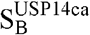 and 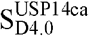) and ten E_D_-like states (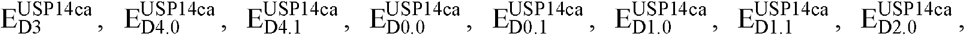, 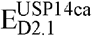 and 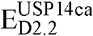). These states were designated by referencing previously published structures^8,9,11,29^ with similar conformational features. Particles with blurred or incomplete densities of certain peripheral subunits (such as RPN1, RPN2 or RPN6) were discarded, provided that the density quality of critical regions of interest (such as the AAA-ATPase ring, the substrate within the central channel, and the ubiquitin tethered to RPN11 or USP14^C114A^) remained unaffected. For the S_D_-like and E_D_-like states, only particles with clear, unblurred density of the USP^C114A^ domain were selected for the final reconstruction, representing the proteasomes that are tightly bound to USP14^C114A^. The resolutions of the final maps were measured by the gold-standard Fourier shell correlation (FSC) in RELION^51^ with the USP14^C114A^-RP subcomplex masked, ranging from 3.6 to 4.5 Å (Extended Data Fig. 3). The 92,349 particles in state 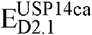 were subjected to another round of focused 3D classification with a mask covering the region around RPN11 and USP14^C114A^, yielding a subclass composed of 16,652 particles showing the substrate density protruding from the entrance of the OB ring to USP14^C114A^ (Fig. 1a,b). Beyond the 16 states, there were additional low-resolution classes among the discarded particles during data processing, with their RP resolution worse than 6 Å. These states were not included in the final reconstruction because their conformational state could not be unambiguously determined due to the poor density of substrates or ATPases.

### Atomic model building and refinement

Using the published coordinates of the USP14-proteasome complex^29^ as initial references, we built atomic models for all 16 conformational states from the USP14^C114A^-proteasome dataset. For each reconstructed cryo-EM map, the coordinates of the closet conformation was first entirely docked into the map and then rigid-body fitted chain by chain in Chimera^54^. Models of chain domains with obvious conformational changes, such as the AAA domains of the six RPT subunits, were manually rigid-body fitted block by block in Coot^55^. Models of all subunits were compared with predicted structures from AlphaFold^56^, and regions with mismatched backbone traces were revised based on the densities. The catalytic Cys114 residue of USP14^C114A^ in each model was mutated to Ala, and the zinc ion at the RPN11 active site was removed if its density is invisible due to the addition of the zinc chelator *o*PA. Subsequent model refinement, including real-space refinement in Phenix^53^ and manual adjustment in Coot^55^, was iterated until the model quality met the expected level.

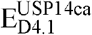 is the only state modeled with specific substrate sequences. In this state, the density of substrate segment within the central channel of the AAA-ATPase ring exhibits several long, bulky side-chain densities (Extended Data Fig. 5p), allowing us to determine the most plausible substrate sequence. Given that this substrate density is immediately below the RPN11-bound ubiquitin, we selected the best-fitting one by fitting all lysine-containing segments of the actual sequence into these densities. The other substrate, observed between the OB ring and USP14^C114A^ but not within the AAA-ATPase channel, is fitted with the N-terminus of the actual sequence based on its length. In all other states, substrates within the AAA-ATPase channel do not show any side-chain densities and thus modelled as repeated polypeptide chains, except for ubiquitylated lysine residues. For states 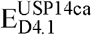 and 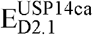, substrate fragments extending from the entrance of the OB ring to USP14^C114A^ were modeled using maps that had been low-pass filtered to 6 Å.

The nucleotide-binding states of ATPases were determined by unbiased modelling of ATP or ADP molecules based on the map densities in the nucleotide-binding pockets and classified as the ‘apo-like’ state if the densities were weak and insufficient to accommodate a nucleotide molecule. Magnesium ions alongside ATP or ADP were assigned based on the map densities only for those maps with the RP resolution better than 3.8 Å (E_A1_, 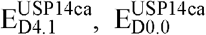 and 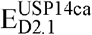).

### Structural analysis and visualization

Structural analysis and visualization were performed in ChimeraX^57^. Comparisons to protein structures from previous publications used the atomic models in the PDB under accession codes: 6EF3 (substrate-engaged Rpn11-inhibited yeast proteasome), 5GJQ (USP14-UbAl-bound human proteasome), 6MSD (state E_A2_ of the USP14-free human proteasome in the act of substrate degradation), 6MSE (state E_B_), 6MSG (state E_C1_), 7W37 (state 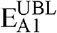 of the USP14^WT^-bound human proteasome in the act of substrate degradation), 7W39 (state 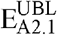), 7W3A (state 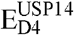), 7W3C (state 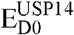), 7W3F (state 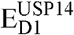), 7W3G (state 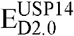), 7W3H (state 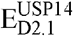), 7W3I (state 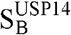) and 7W3J (state 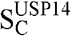).

### Conformational landscape analysis of the time-resolved cryo-EM dataset

To visualize conformational changes over time more intuitively, we reconstructed conformational landscapes of the time-resolved dataset alongside previously published data (Extended Data Fig. 12a-c). Particles from this dataset (464,682 in total) and the previous study^29^ (1,194,524 with ATPγS and 1,494,793 without ATPγS) were combined and aligned to a common reference using CP-masked auto-refinement in RELION^51^. The combined dataset was then randomly divided into 20 subsets, each subjected to alignment-skipped RP-masked 3D classification, with the class number set to 20. A total of 400 class averages were generated. Each class was independently reconstructed in RELION^51^, and the resulting map was multiplied by a common RP mask and low-pass filtered to 6 Å. The conformational variability of these maps was analyzed and embedded into a 2D conformational landscape using deep residual autoencoder and *t*-distributed stochastic neighbor embedding (t-SNE) in AlphaCryo4D^52^. This landscape provides a representation of the conformational distributions of all particles. The free energy value of each point in the landscape was calculated as:

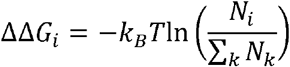

where ΔΔ*G*_*i*_ denotes the free energy difference of a data point with the particle number of *N*_*i*_ against a common reference energy level, *k*_*B*_ is the Boltzmann constant and *T* is the temperature in Kelvin. By substituting the corresponding particle numbers, we generated landscapes in the same t-SNE coordinate system for different datasets or time points (Fig. 5 and Extended Data Fig. 12d-g). This allowed us to compare the structural heterogeneity under various biochemical conditions and track changes of the conformational space over time.

## Author Contributions

S.Zou, S.Zhang and Y.M. conceived this study. S.Zhang designed and purified the substrates for degradation assays. S.Zou and L.Z. purified the other proteins and conducted the biochemical experiments. S.Zou and S.Zhang prepared cryo-EM samples, collected cryo-EM data and analyzed the experimental cryo-EM datasets. S. Zhang refined the density maps, built the atomic models and analyzed the data. S.Zhang, S.Zou and Y.M. wrote the manuscript. Y.M. supervised the project.

## Acknowledgements

We thank Y. Saeki for the plasmids expressing Sic1^PY^ and WW-HECT; L. Huang for the proteasome-expressing cell line; C. Tang for the plasmid expressing UBE1; X. Li, Z. Guo, X. Pei, C. Fan and Y. Ma for technical assistance with cryo-EM data collection and storage; D. Liu and Q. Zhang for technical assistance with the mass spectrometry experiments. The cryo-EM data were collected at the Cryo-EM Core Facility Platform and Laboratory of Electron Microscopy at Peking University. The data processing was supported by the High-Performance Computing Platform of Peking University. The mass spectrometry data were collected at the National Center for Protein Sciences at Peking University. This work was supported by National Key Research and Development Program of China (2023YFF1204400 and 2023YFF1204401 to Y.M.), National Natural Science Foundation of China (12125401, 12090051 and 11774012 to Y.M. and 32471308 to S.Zou), Beijing Natural Science Foundation grant (Z180016/Z18J008 to Y.M.), and China Postdoctoral Science Foundation (2023T160003 to S.Zhang).

## Data availability

Atomic coordinates and cryo-EM structures have been deposited in the Protein Data Bank (and the EMDB) with accession codes 9JY5 (EMD-61888), 9JY6 (EMD-61889), 9JY7 (EMD-61890), 9JY8 (EMD-61891), 9JY9 (EMD-61892), 9JYA (EMD-61893), 9JYB (EMD-61894), 9JYC (EMD-61895), 9JYD (EMD-61896), 9JYE (EMD-61897), 9JYF (EMD-61898), 9JYG (EMD-61899), 9JYH (EMD-61900), 9JYI (EMD-61901), 9JYJ (EMD-61902), 9JYK (EMD-61903). All raw data are available upon reasonable request.

## Competing interests

The authors declare no competing interests.

---

**Extended Data Fig. 1:**
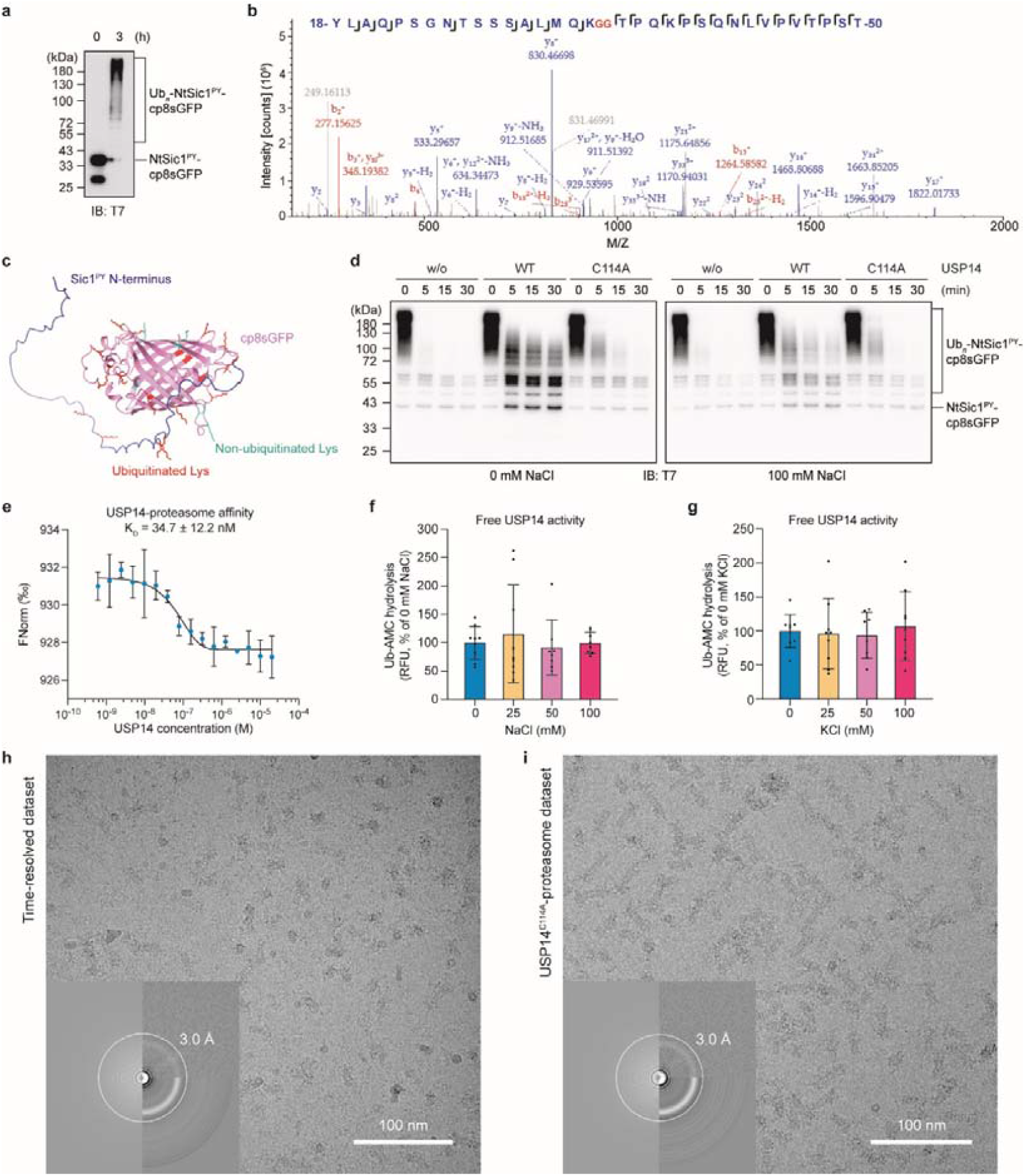
Purification and biochemical characterization of the newly engineered model substrate, and activity assays and cryo-EM imaging of USP14 and the proteasome. **a, b**, Ubiquitylation of the substrate NtSic1^PY^-cp8sGFP, analyzed by SDS-PAGE and western blot using an anti-T7 antibody (**a**), and by LC-MS/MS analysis (**b**). A representative tandem mass spectrum from the LC-MS/MS assay (**b**) shows ubiquitylation at Lys34. **c**, The predicted atomic model of NtSic1^PY^-cp8sGFP, generated by AlphaFold^56^. Ubiquitylated (red) and non-ubiquitylated (cyan) lysine residues identified by the LC-MS/MS assay are highlighted, with their side chains displayed in stick representation. **d**, *In vitro* degradation assay of Ub_*n*_-NtSic1^PY^-cp8sGFP at 30 °C, performed in the presence or absence of USP14 variants at different NaCl concentrations, analyzed by SDS-PAGE and western blot using an anti-T7 antibody. This assay validates the results shown in Fig. 4e. **e**, Microscale thermophoresis (MST) analysis of USP14 binding to the human proteasome in the presence of 100 mM NaCl, with the dissociation constants (*K*_D_) calculated from three independent experiments (shown as mean ± s.d.). As reported in our previous study^29^, the K_D_ between USP14 and the proteasome in the absence of NaCl is 94.6□±□27.1□nM, which is higher than the corresponding K_D_ in the presence of 100 mM NaCl. **f, g**, Ubiquitin-AMC hydrolysis by USP14 in the absence of the proteasome under varying NaCl (**f**) or KCl (**g**) concentrations. Data are presented as mean ± s.d. from three independent experiments, each consisting of three replicates. RFU, relative fluorescence units. **h, i**, Typical motion-corrected cryo-EM micrographs of the substrate-engaged human USP14^WT^-proteasome complex from the time-resolved dataset at a reaction time point of 1 minute (**h**), alongside the RPN11-inhibited USP14^C114A^-proteasome in the presence of substrates (**i**). The lower left insets of both panels show the power spectrum evaluations of the corresponding micrographs, analyzed using GCTF^48^.

**Extended Data Fig. 2:**
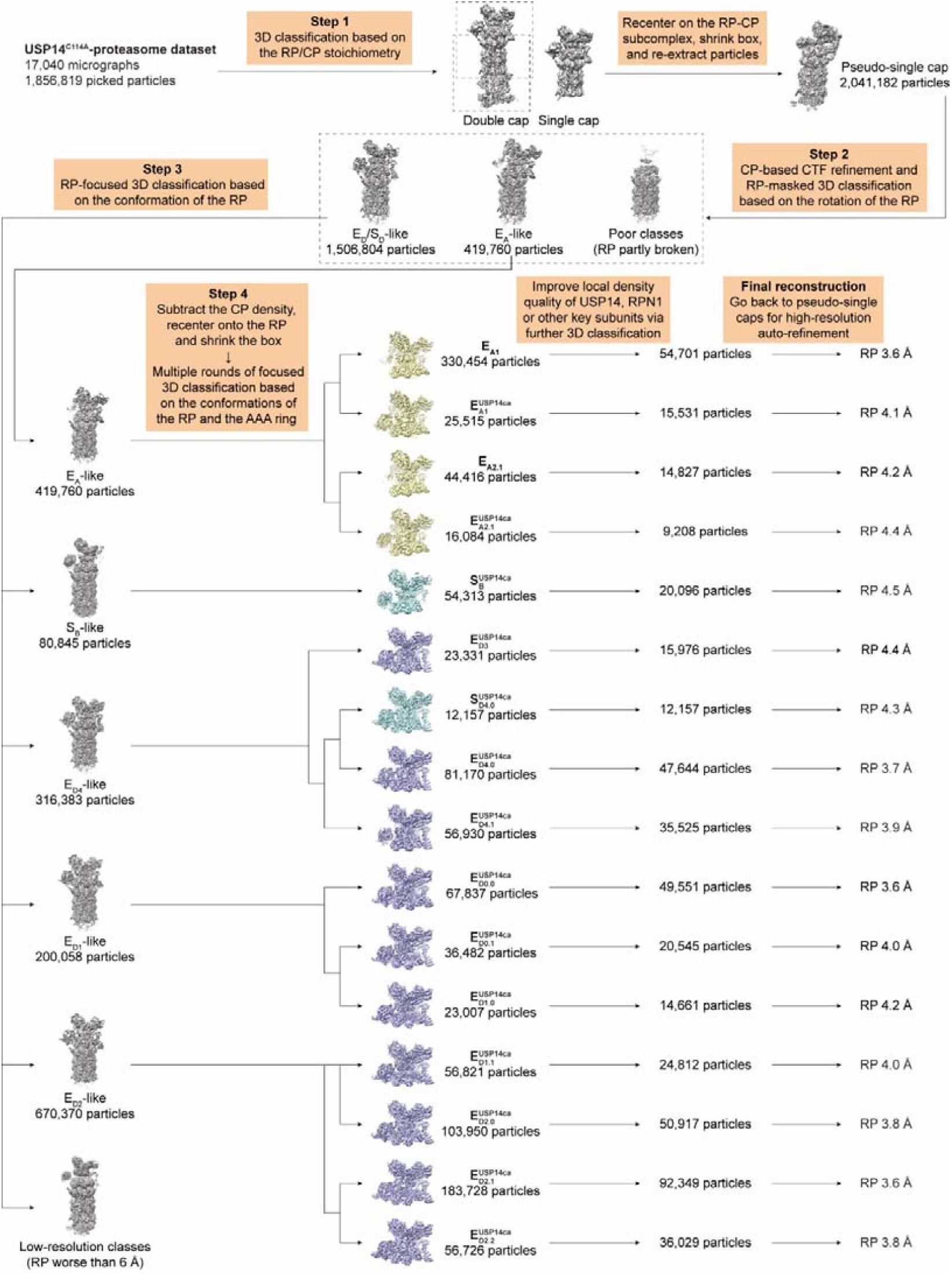
Cryo-EM data processing for the RPN11-inhibited USP14^C114A^-proteasome complex. The flowchart illustrates the major steps of the hierarchical 3D classification strategy. Detailed iterations of classification in each step are omitted for clarity. After step 4, the maps of the E_A_-like, S_D_-like and E_D_-like states are colored yellow, cyan and violet, respectively. The particle numbers after 3D classification and final reconstruction for each state, as well as the resolutions of the RP-masked reconstructions, are labelled in the figure.

**Extended Data Fig. 3:**
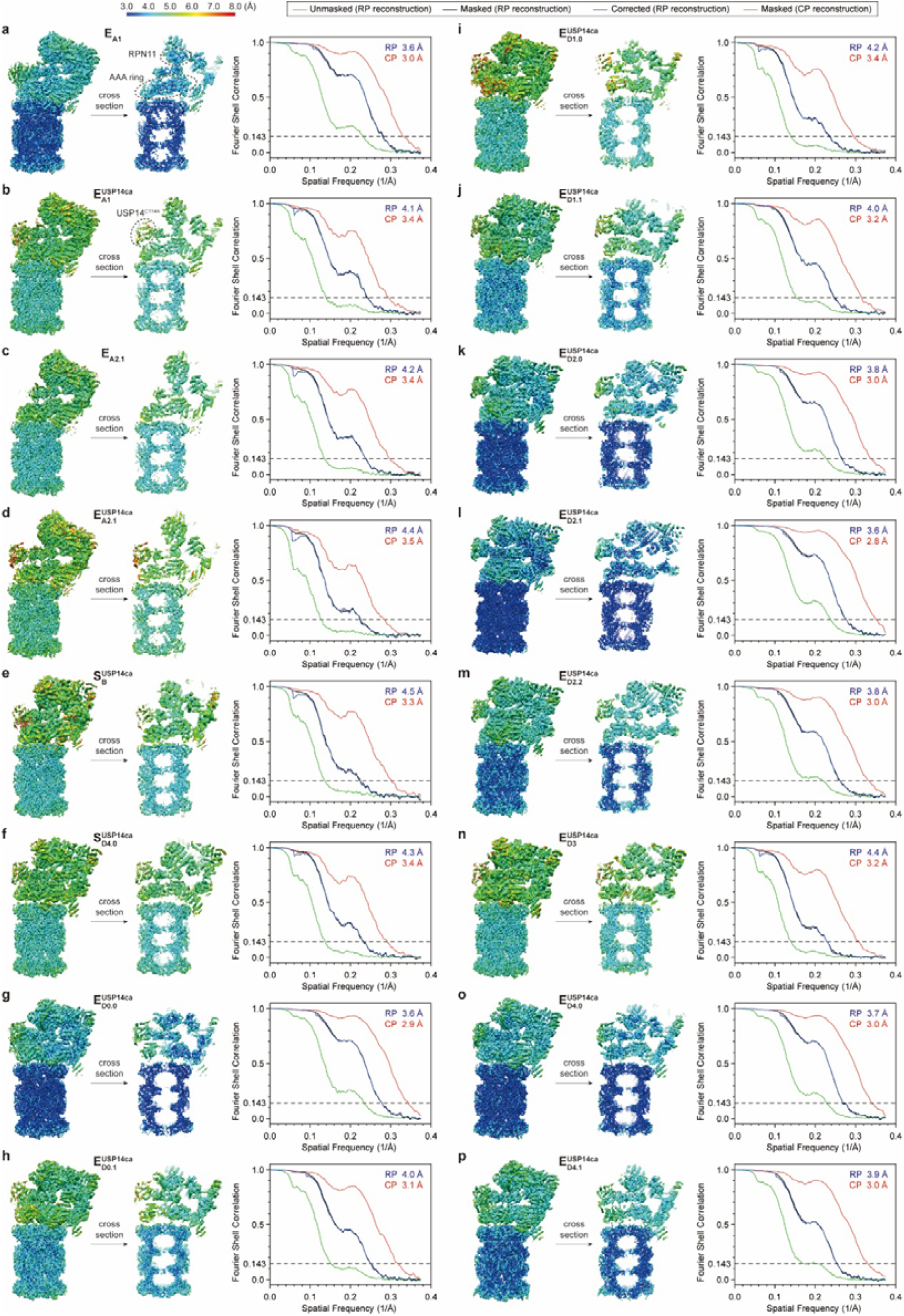
Cryo-EM reconstructions and resolution measurements of the RPN11-inhibited USP14^C114A^-proteasome complex. Each panel presents one state and is organized from left to right in the same order as follows: a side view (left) and its cross section (middle) of the complete RP-CP map colored based on local resolution, and gold-standard Fourier shell correlation (FSC) plots labelled with estimated resolutions for both RP-masked and CP-masked reconstructions (right). The local resolution maps and the FSC curves were calculated using ResMap^58^ and RELION^51^, respectively. A consistent color bar for all local resolutions is displayed in the upper-left inset. For the RP-masked reconstruction of each state, the figure shows the unmasked (green), masked (black) and corrected (blue) FSC curves. For the CP-masked reconstruction of each state, only masked FSC curve (red) is presented.

**Extended Data Fig. 4:**
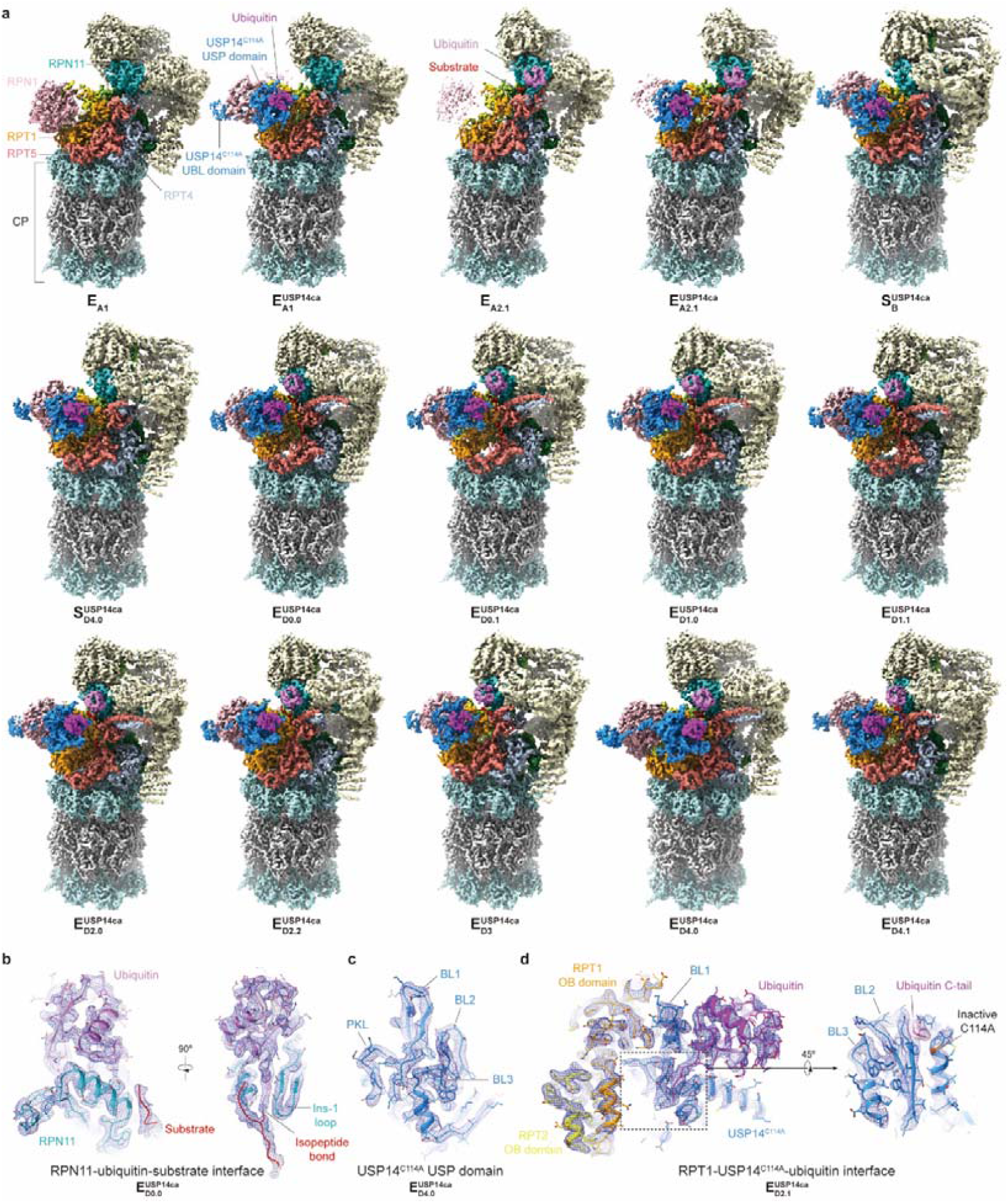
Cryo-EM maps of the RPN11-inhibited USP14^C114A^-proteasome complex. **a**, Gallery of refined cryo-EM maps of all states except for 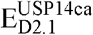, which is shown in Fig. 1a. **b**-**d**, High-resolution cryo-EM densities (mesh) of the RPN11-ubiquitin-substrate interface in state 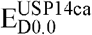 (**b**), the USP^C114A^ domain in state 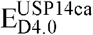 **(c)** and the RPT1-USP14 -ubiquitin interface in state 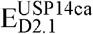 (**d**), each superimposed with their atomic models. Critical loops of the USP^C114A^ domain are labelled in panels **c** and **d**. The C114A mutation is highlighted in orange in the right image of panel **d**.

**Extended Data Fig. 5:**
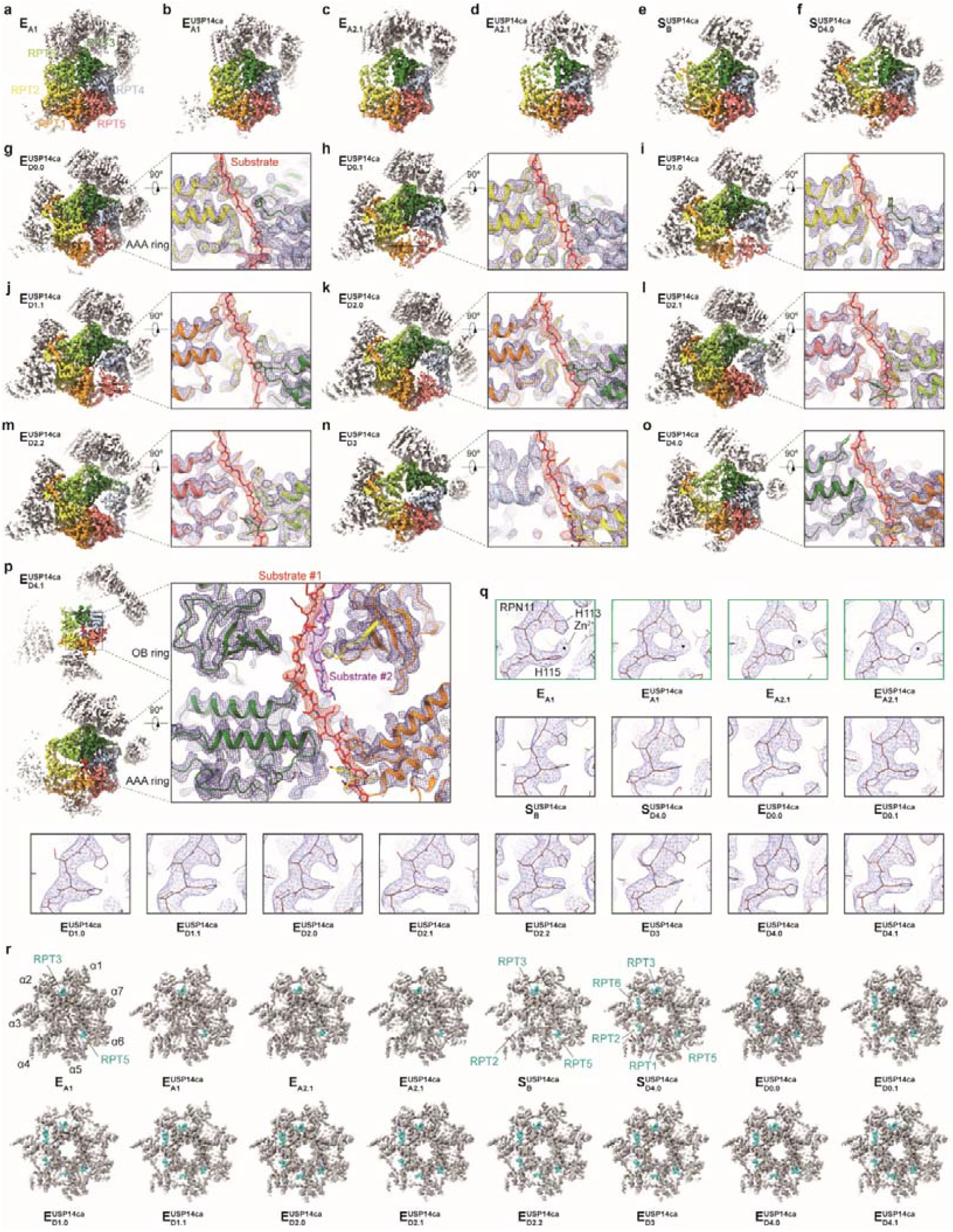
Key structural features of the RPN11-inhibited USP14^C114A^-proteasome in different states. **a**-**p**, Cryo-EM densities of the proteasomal AAA-ATPase ring in different states. Panels **a**-**f** and the left insets of panels **g**-**p** show top views of the AAA-ATPase ring. Panel **p** also includes a top view of the OB ring (upper left) in state 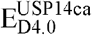. The right insets of panels **g**-**p** show zoomed-in side views of the substrate (red) interacting with the ATPase pore-1 loops, shown in mesh representation. The substrates in panels **g**-**p** are modelled as polypeptide backbones and shown as sticks. Models of the AAA-ATPase ring and side chains of the aromatic residues of the pore-1 loops are shown as cartoon and sticks, respectively. **q**, Cryo-EM densities (mesh) and fitted models (stick) of the RPN11 active site in different states. The catalytic zinc ion is visible and marked by a dot in the four E_A_-like states. **r**, Cryo-EM densities of the top views of the RP-CP interface in different states. The C-terminal tails of the RPT subunits inserted into the α-pockets of the CP are colored in cyan. The CP gate remains closed in the four E_A_-like states and in state 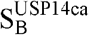 but is fully open in the other states.

**Extended Data Fig. 6:**
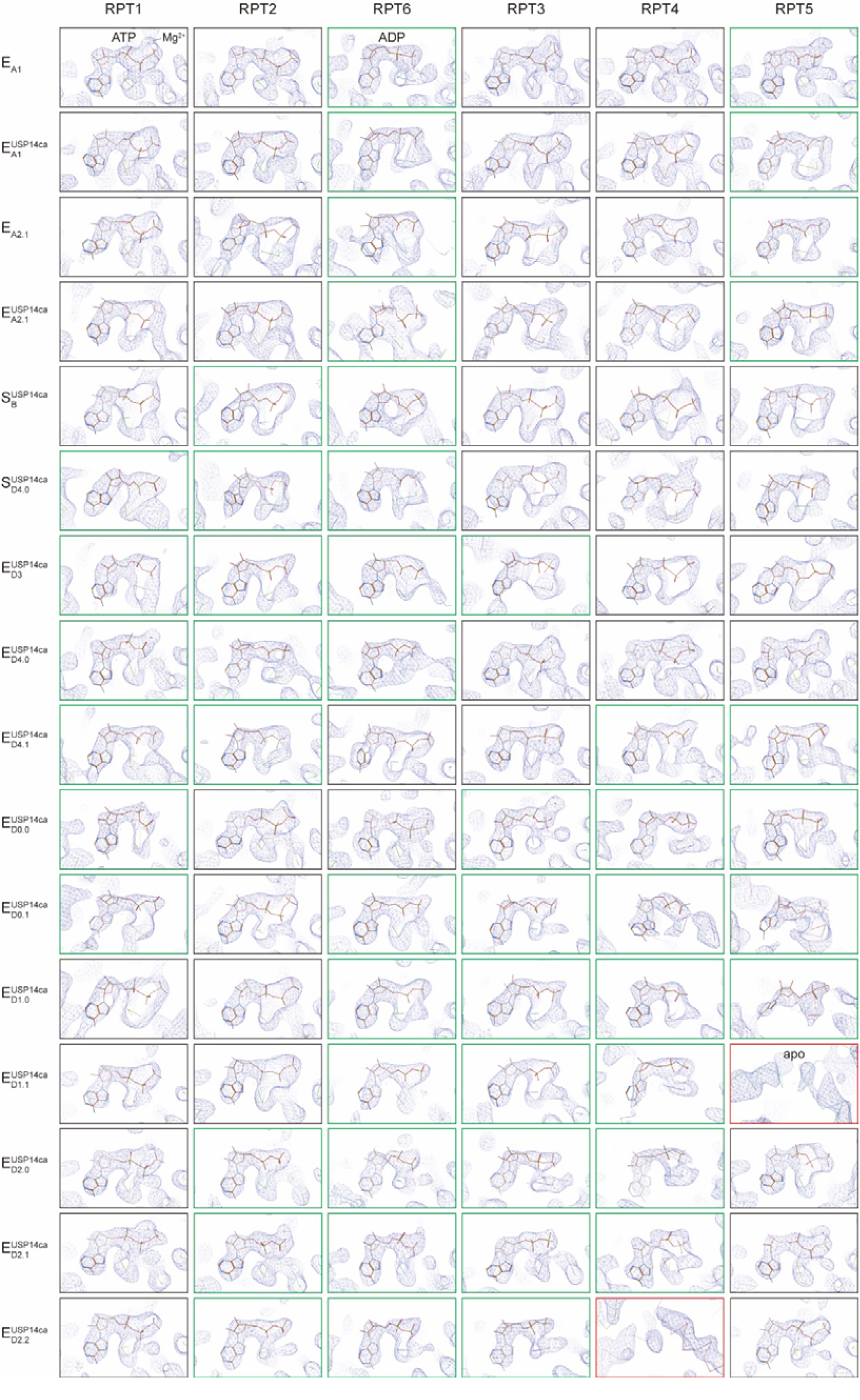
Nucleotide-binding states of the six ATPases in the RPN11-inhibited USP14^C114A^-proteasome across different states. All close-up views are direct screenshots of the ATPase nucleotide-binding pockets in Coot^55^ after atomic modelling into the density maps (mesh), presented without modification. Boxes displaying ATP-bound, ADP-bound and apo-like subunits are outlined in black, green and red edges, respectively. At the contour level typically used for atomic modelling, the potential nucleotide densities in the apo-like subunits largely disappear, although they can occasionally appear as partial nucleotide shapes at much lower contour levels. For maps with the RP resolution better than 3.8 Å (E_A1_, 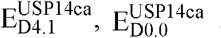 and 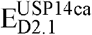), magnesium ions associated with ATP or ADP were assigned based on the density maps.

**Extended Data Fig. 7:**
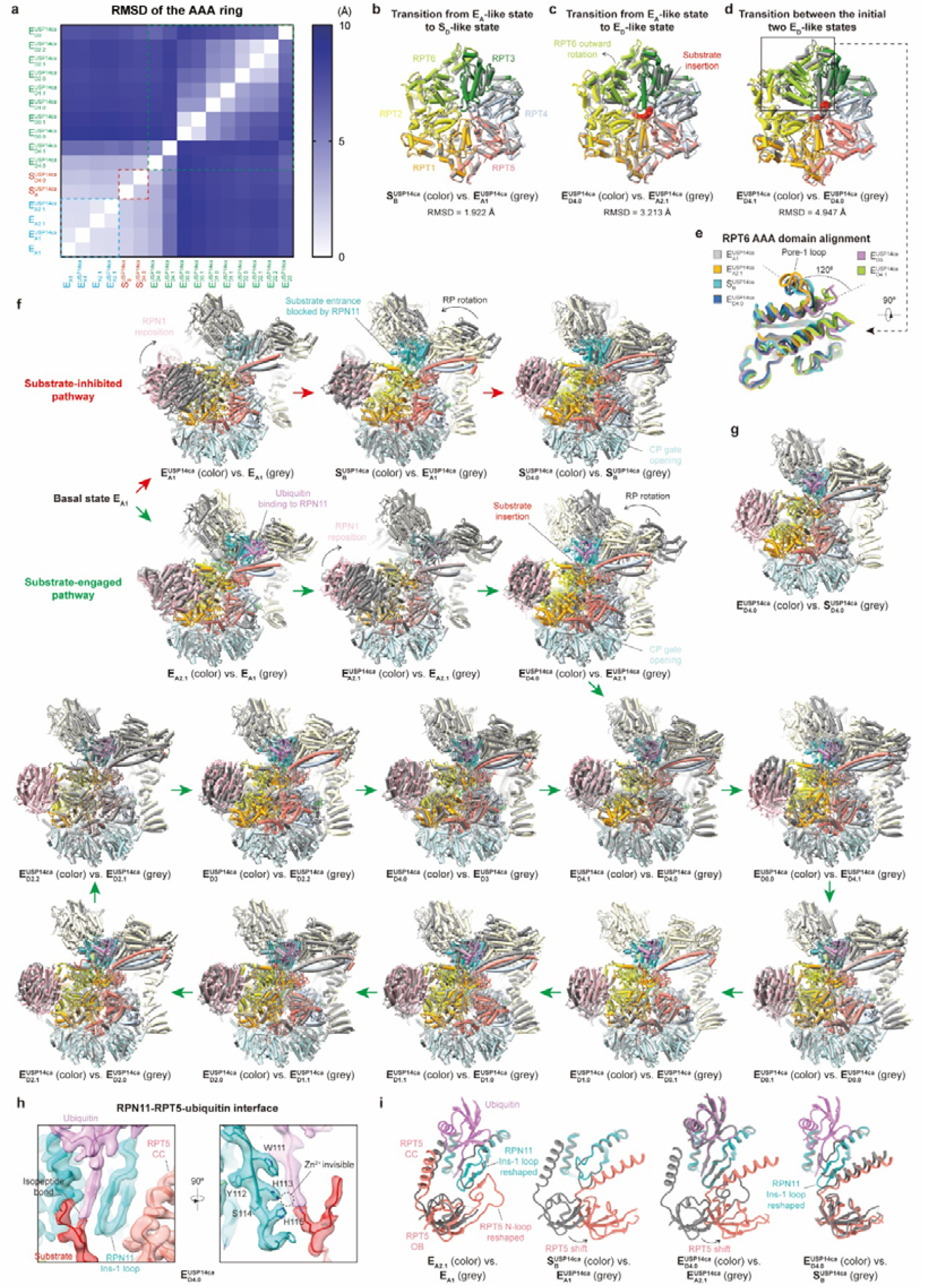
Structural comparison of the RPN11-inhibited USP14^C114A^-proteasome across different states. **a**, Root-mean-squared-deviation (RMSD) of the AAA-ATPase ring structures mapped between pairs of the 16 states. **b**-**d**, Structural comparison of the AAA-ATPase ring related to key state transitions: from an E_A_-like state to an S_D_-like state (**b**), from an E_A_-like state to an E_D_-like state following substrate insertion (**c**) and between the initial two E_D_-like states (**d**). The models of substrates are shown in sphere representation in panels **c** and **d. e**, Structural comparison of the RPT6 AAA domain across all states shown in panels **b**-**d** as well as state 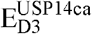. The models are superimposed after alignment of the RPT6 large AAA subdomain. States 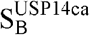 and 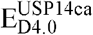 exhibit similar conformations of the RPT6 AAA domain to those seen in the E_A_-like states. **f**, Structural comparison of the RP between adjacent states, step by step, along two state-transition pathways. The models of the CP are used for alignment and are partially hidden for clarity. The models of USP14^C114A^ are also hidden to better illustrate the conformational changes in the RP. Red and green arrows indicate substrate-inhibited and substrate-engaged pathways, respectively. **g**, Structural comparison of the RP between the substrate-engaged state 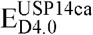 and substrate-free state 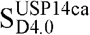. These two states display similar RP conformations, except for the difference being the presence of substrate in 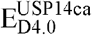. **h**, Close-up view of the uncleaved isopeptide bond linking the substrate lysine and the ubiquitin C-terminus bound in the catalytic groove of RPN11 in state 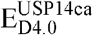. The catalytic zinc ion is not visible due to *o*PA treatment. The Ins-1 loop of RPN11 forms a β-hairpin, sandwiched between the ubiquitin C-terminus and the RPT5 CC domain. **i**, Structural comparison of the RPN11-RPT5-ubiquitin interface between pairs of representative states. The models are superimposed after alignment of the RPN11 subunit. The Ins-1 loop of RPN11 adopts the β-hairpin conformation in states E_A2.1_, 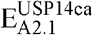 and all E_D_-like states. RPT5 is repositioned relative to RPN11 in all S_D_-like and E_D_-like states.

**Extended Data Fig. 8:**
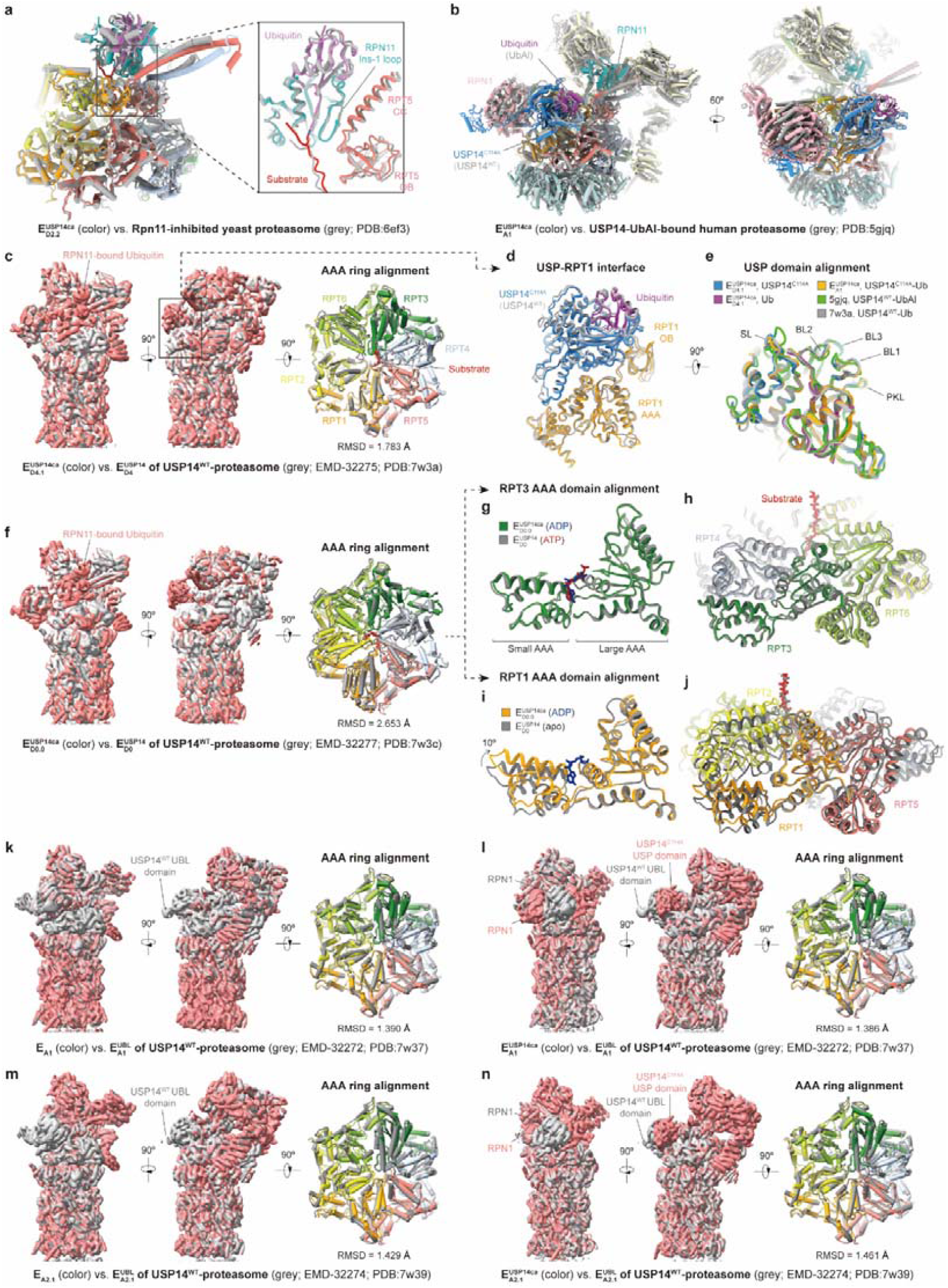
Structural comparison of the RPN11-inhibited USP14^C114A^-proteasome with previously reported cryo-EM structures. **a**, Structural comparison of the RPN11-RPT subcomplex in state 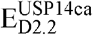 with a cryo-EM structure of the substrate-engaged Rpn11-inhibited yeast proteasome in a comparable conformation^36^ (PDB ID: 6EF3). The two models are superimposed after alignment using the AAA-ATPase ring (left) and RPN11 (right). **b**, Structural comparison of the USP14-RP subcomplex in state 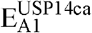 with the cryo-EM structure of the USP14-UbAl-bound human proteasome^27^ (PDB ID: 5GJQ). The models of the CP are used for alignment and are partially hidden for visual clarity. **c**-**n**, Structural comparison of the 26S proteasome holoenzyme and the AAA-ATPase ring across different states with previously determined cryo-EM structures of the USP14^WT^-bound proteasome in comparable states^29^. For panels **c, f** and **k**-**n**, the images on the left and middle show superimposed density maps of the proteasome, low-pass filtered to 8 Å, and the images on the right show superimposed models of the AAA-ATPase ring. **d**, Close-up view of the USP14-ubiquitin-RPT1 models of the boxed region in panel **c. e**, Superimposed models of the ubiquitin-bound USP domain of all structures shown in panels **b** and **c**. The USP^C114A^ domain and ubiquitin in state 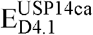 are individually colored for visual differentiation, with critical loops of the USP domain labelled. **g**-**j**, Structural comparison of the ATPases between states 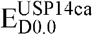 (color) and 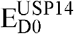 (grey; PDB ID: 7W3C). Atomic models of the individual subunits (**g, i**) and the entire AAA-ATPase ring (**h, j**) are shown after alignment of the large AAA subdomain of RPT3 (**g, h**) or RPT1 (**i, j**).

**Extended Data Fig. 9:**
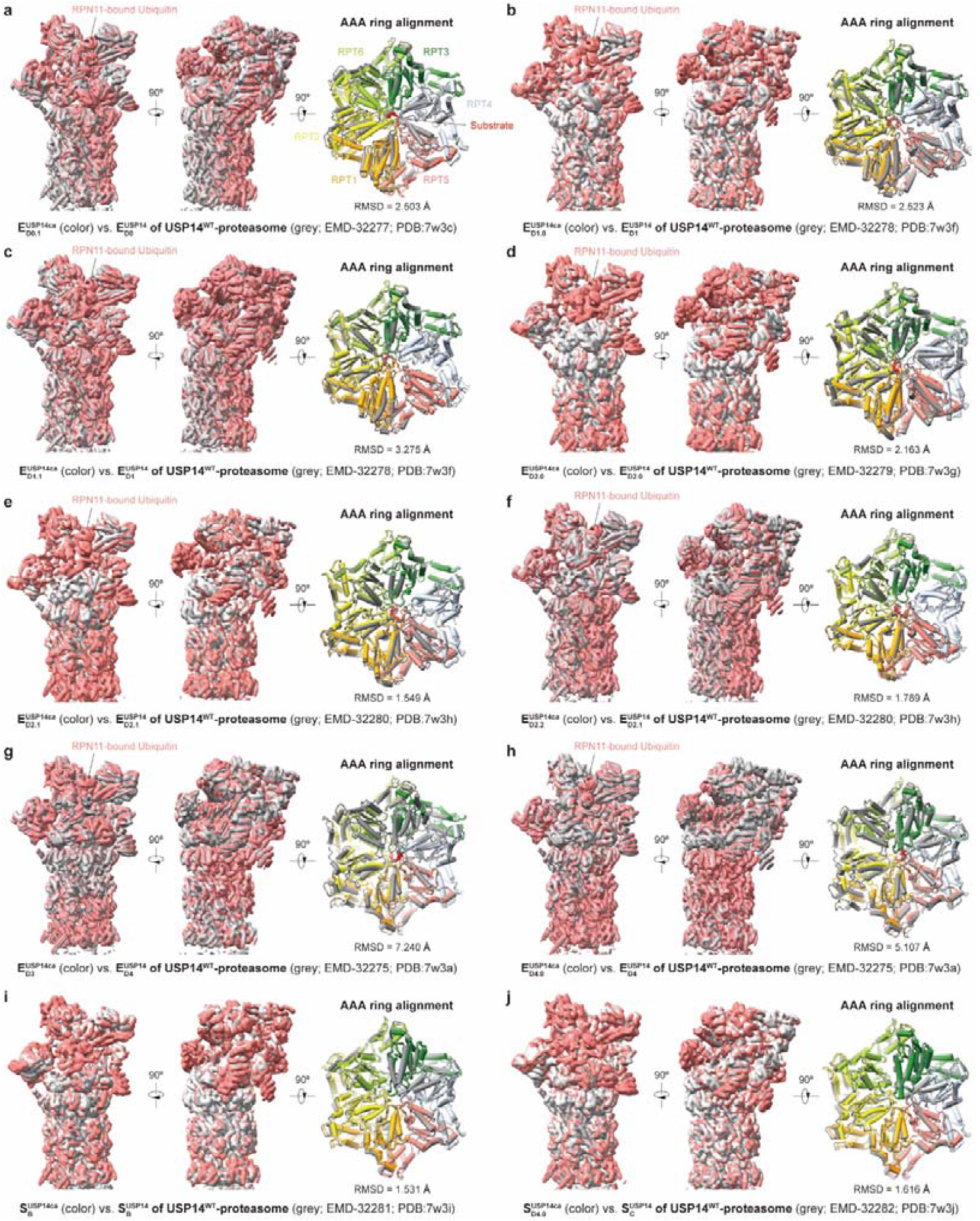
Structural comparison of the RPN11-inhibited USP14^C114A^-proteasome with the USP14^WT^-proteasome. The figure includes comparisons of states that are not displayed in Extended Data Fig. 8c-n. In each panel, the images on the left and middle show superimposed density maps of the 26S holoenzyme, low-pass filtered to 8 Å, and the images on the right show superimposed models of the AAA-ATPase ring.

**Extended Data Fig. 10:**
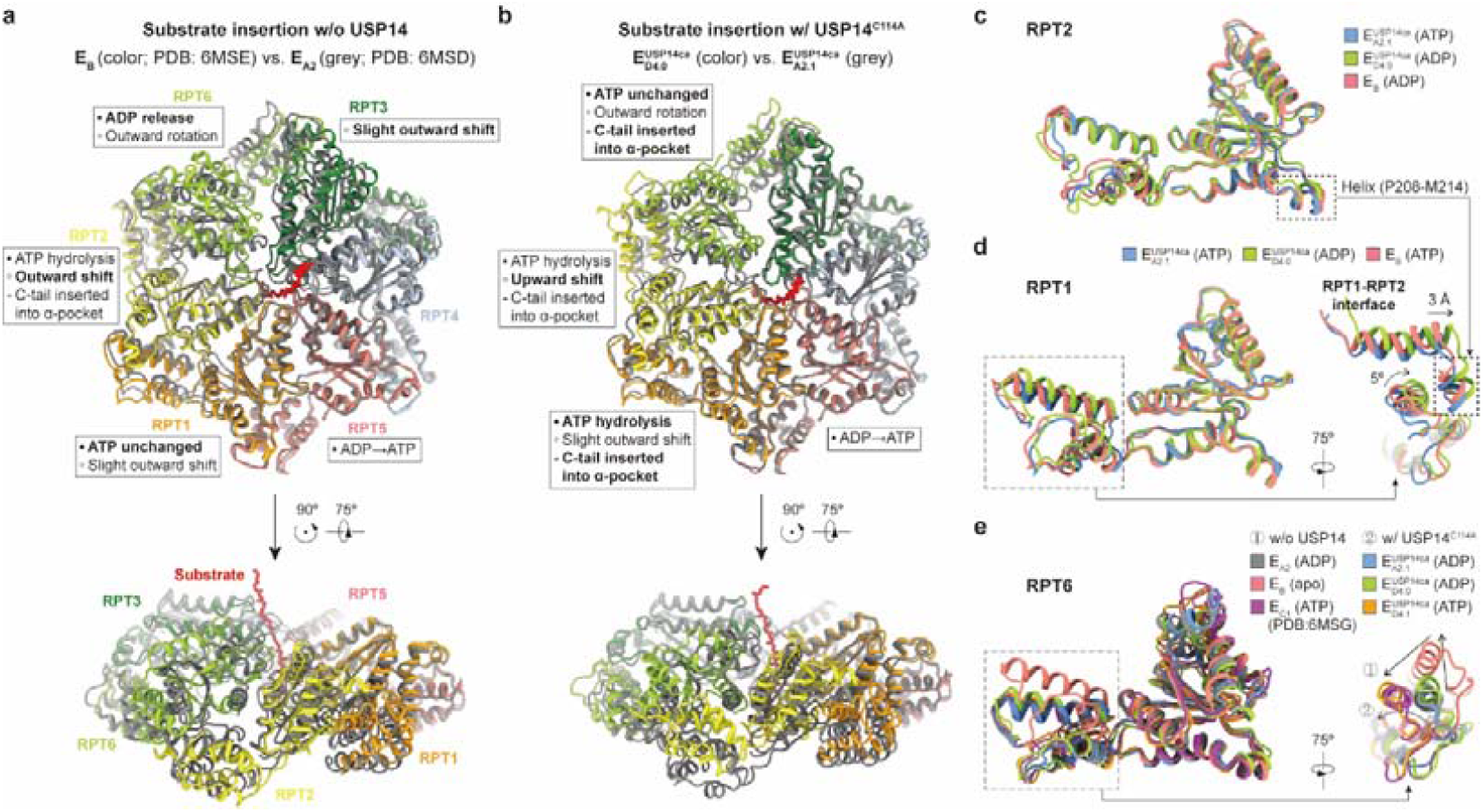
Conformational changes in the AAA-ATPase ring induced by substrate insertion into the central channel, comparing conditions with and without USP14^C114A^. **a, b**, Overall conformational changes of the AAA-ATPase ring triggered by substrate insertion, observed in the absence of USP14 **(a)** and in the presence of USP14^C114A^ (**b**). The models are superimposed after alignment of the AAA-ATPase ring, shown in both top view (upper) and tilt view (lower). Conformational changes of each ATPase subunit are annotated, including the nucleotide-binding state, overall motion and the insertion status of the C-terminal tail. These changes are differentiated with distinct bullet points, with notable changes emphasized in bold. **c**-**e**, Superpositions of the AAA domain structures of RPT2 (**c**), RPT1 (**d**) or RPT6 (**e**) in distinct states associated with substrate insertion or translocation initiation, aligned based on their large AAA subdomains. The models of ATP and ADP are hidden for clarity. The short helix of RPT2 from Pro208 to Met214 is also depicted in panel (**d**) to emphasize changes at the RPT1-RPT2 interface. In the right image of panel (**e**), black arrows indicate the motion directions of the RPT6 small AAA subdomain in the absence or presence of USP14^C114A^.

**Extended Data Fig. 11:**
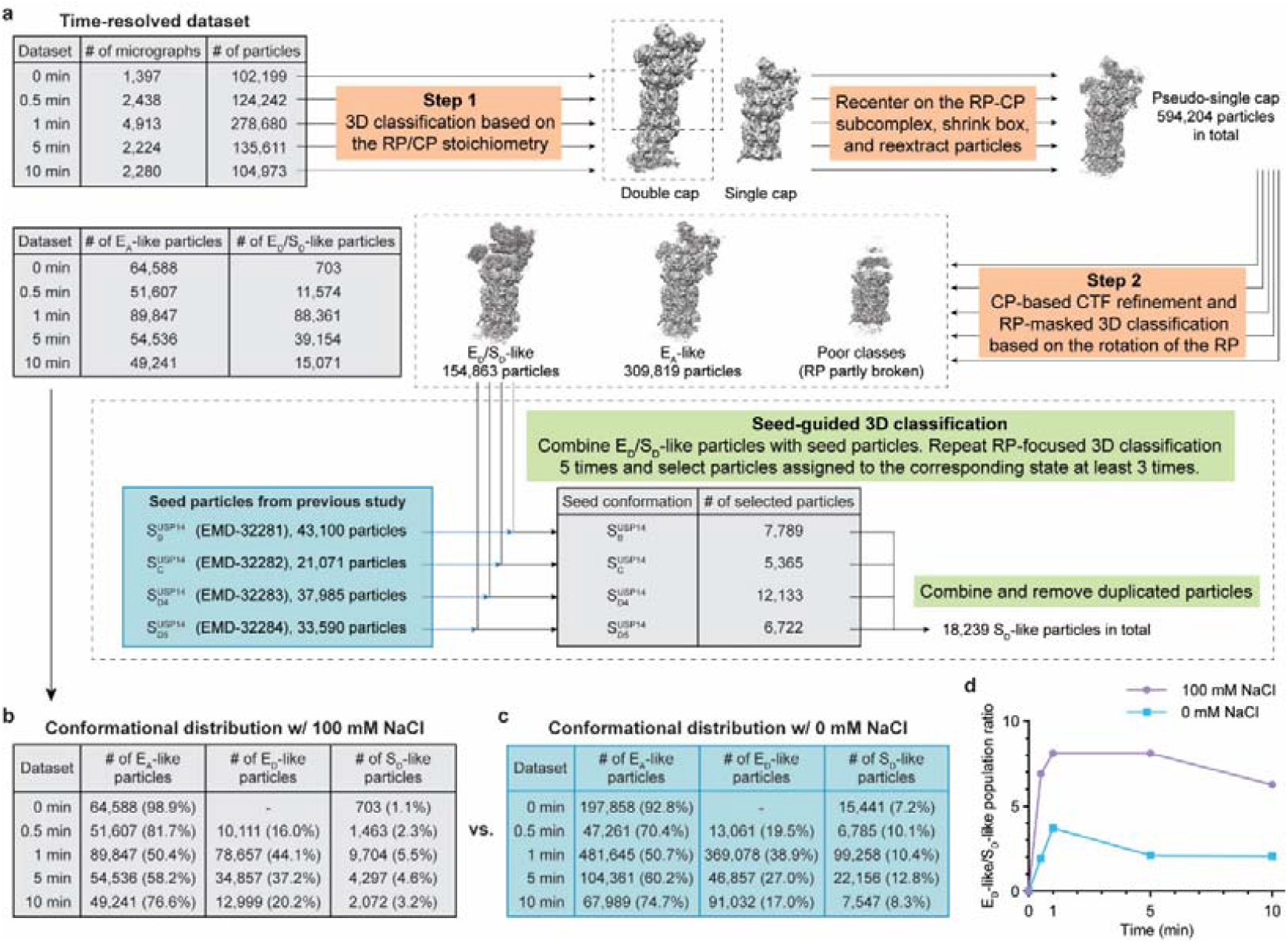
Data processing of the time-resolved cryo-EM datasets. **a**, Flowchart of the major steps of the hierarchical 3D classification strategy. Detailed iterations of classification in each step are omitted for clarity. The blue box represents proteasome particles in S_D_-like states from our previous time-resolved cryo-EM study^29^, which were used as seeds for seed-guided 3D classification. The analysis is not aimed at achieving high-resolution reconstructions but rather for a fair comparison with previous studies. **b, c**, Time-resolved conformational distribution of the three major states in the presence of 100 mM NaCl (**b**) or in the absence of NaCl (**c**). Data in panel (**c**) are sourced from the previous study^29^. The histograms based on panels (**b**) and (**c**) are shown in Fig. 4b. **d**, Kinetic changes in the ratio of E_D_-like to S_D_-like state populations based on the data presented in panels **b** and **c**.

**Extended Data Fig. 12:**
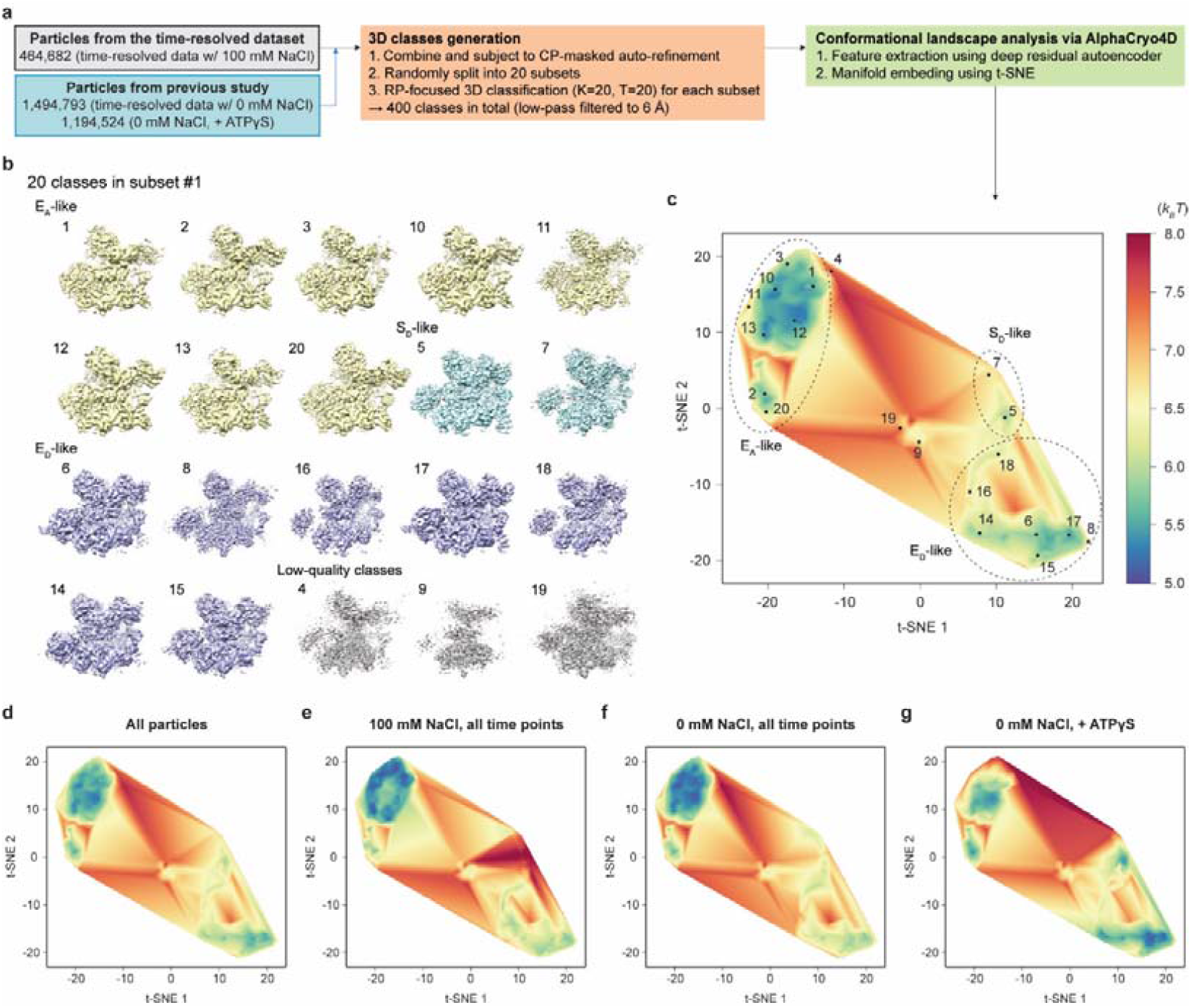
Conformational landscape analysis of the time-resolved datasets. **a**, Workflow for the conformational landscape analysis of the time-resolved dataset in combination with data from the previously published study^29^. **b**, RP-mask 3D Classification for a typical subset that results in 20 maps. The maps are all low-pass filtered to 6 Å. Three major categories of states and low-quality classes are distinguished with different colors. **c**, Conformational landscape of all particles listed in panel **a**. The dots indicate the locations on the landscape corresponding to the 20 maps shown in panel **b**. The dashed ellipses mark the regions occupied by three major categories of states. **d**-**g**, Conformational landscapes of the USP14-proteasome separated according to different conditions: all particles (**d**), in the presence of 100 mM NaCl (**e**), in the absence of NaCl (**f**) and with the addition of ATPγS (**g**). The same t-SNE coordinate system and the color bar shown in panel **c** are used for all landscapes in panels **c**-**g** and those in Fig. 5 of the main text. Datasets for panels **f** and **g** are from the previous study^29^.

## Notes

### Competing Interest Statement

The authors have declared no competing interest.

